# Cardiomyocyte-specific plakophilin-2 loss is sufficient to induce aging and senescence of nonmyocytes. Relevance to arrhythmogenic cardiomyopathy

**DOI:** 10.1101/2025.10.22.682160

**Authors:** Giorgia Bertoli, Kavya Phadke, Alessandro Cospito, Joanna Abi Rizk, Mingliang Zhang, Eleni Miliotou, Michael Cammer, Yan Deng, Valeria Mezzano, Mark Alu, Gyles Ward, Cynthia Loomis, Adriana Heguy, Feng-Xia Liang, Eric M. Small, Irene de Lázaro, Mario Delmar

**Affiliations:** The Leon H. Charney Division of Cardiology, New York University Grossman School of Medicine, New York, NY, USA; Department of Biomedical Engineering, NYU Tandon School of Engineering, New York University, New York, NY, USA; Microscopy Laboratory [RRID:SCR_017934], Division of Advanced Research Technologies, New York University Grossman School of Medicine, New York, NY, USA; Experimental Pathology Research Laboratory [RRID:SCR_017928], Division of Advanced Research Technologies, NYU Grossman School of Medicine, New York, NY, USA; The Genome Technology Center [RRID:SCR_017929], New York University Grossman School of Medicine, New York, NY, USA; Aab Cardiovascular Research Institute, Department of Medicine, University of Rochester School of Medicine and Dentistry, Rochester, NY, USA

**Keywords:** Arrhythmias, Cardiomyopathies, Sudden Death, Inflammation, Biological age

## Abstract

**Introduction:** Pathogenic variants in *PKP2* are the most common cause of familial arrhythmogenic right ventricular cardiomyopathy (ARVC).

**Objective:** To test whether PKP2 deficiency only in cardiomyocytes is sufficient to provoke premature aging and pro-inflammatory senescence in non-myocytes, cardiac resident cells.

**Methods:** We studied mice with cardiomyocyte-specific, tamoxifen-activated loss of PKP2 (PKP2cKO) using conventional and multiplex imaging, cytokine arrays, epigenetic clocks, spatial transcriptomics, expansion and structured illumination microscopy, and correlative data analysis. We examined non-myocytes and cardiomyocytes for premature aging and senescence.

**Results:** We observed senescence-associated heterochromatin foci (SAHFs) and p21 staining in non-myocytes. Cytokines in media of non-myocyte cells were consistent with senescence-associated secretory phenotype (SASP). Epigenetic clocks identified premature aging. Multiplex immunohistochemistry showed non-myocyte cells in niches, intermingled with cardiomyocytes. Spatial transcriptomics showed over-representation of SASP-related transcripts, predominantly in myocyte-rich areas of the left ventricle. SAHFs, p21 staining and increased epigenetic age were not found in cardiomyocytes from PKP2cKO hearts, though we observed structural features associated to premature aging. Cross-reference analysis showed correlation between the PKP2cKO cardiac proteome and that of mice 5 or 6 times their chronological age, as well as transcriptional signatures of neurodegenerative diseases.

**Conclusion:** Loss of PKP2 expression only in adult cardiac myocytes is sufficient to induce pro-inflammatory senescence in non-myocytes, and overall premature cardiac aging. This is the first study to intersect cellular senescence and premature aging in desmosomal arrhythmogenic cardiomyopathies. We speculate that cell-agnostic molecular signatures, biomarkers, and pharmacology of senescence and of neurodegenerative diseases may be relevant to diagnose or treat PKP2-ARVC.

**UNSTRUCTURED ABSTRACT:** Pathogenic variants in *PKP2* are the most common cause of familial arrhythmogenic right ventricular cardiomyopathy. We used the PKP2cKO model (cardiomyocyte-specific, tamoxifen-activated PKP2 knockout) to examine whether PKP2 deficiency only in cardiomyocytes is sufficient to provoke premature aging and pro-inflammatory senescence in non-myocyte, cardiac resident cells. Through a variety of methods, we identified senescence-associated heterochromatin foci (SAHFs), p21 staining, and cytokines consistent with senescence-associated secretory phenotype (SASP) in cells and media of non-myocytes. Epigenetic clocks identified premature aging. Spatial transcriptomics showed over-representation of SASP-related transcripts, predominantly in myocyte-rich areas of the left ventricle. SAHFs, p21 staining and epigenetics suggesting advanced age were not found in cardiomyocytes, though we observed structural features associated to premature aging. Cross-reference analysis showed correlation between the PKP2cKO cardiac proteome and that of mice 5 or 6 times their chronological age, as well as transcriptional signatures of neurodegenerative diseases.

**HIGHLIGHTS:**

- PKP2-ARVC is a leading cause of sudden unexpected death in the young.
- The molecular path from the variant of a gene to the clinical disease remains unclear. An inflammatory component has been postulated.
- We show that loss of PKP2 only in myocytes is sufficient to induce a pro-inflammatory senescence in non-myocytes and premature aging of cardiac cells.
- Similarities to the molecular profile of neurodegenerative diseases and novel paths to therapy are discussed.

## INTRODUCTION

Plakophilin-2 (PKP2) is a component of the desmosome and of the cardiac intercalated disc as a whole. ^1^ Pathogenic variants in the *PKP2* gene are the most common cause of familial arrhythmogenic right ventricular cardiomyopathy (PKP2-ARVC), a condition associated with sudden cardiac arrest in young individuals. ^2^ Disease progression can lead to end-stage heart failure and the need for heart transplant. ^2^ As with other conditions associated to gene deficiencies, the steps that connect the defect of a gene to the failure of an organ remain unclear. Extensive research has unveiled deficits in chemical, ^3,4^ metabolic^5–7^ and mechanical function^5^ that directly or indirectly cause changes in the transcriptional program of the cells of hearts with a desmosomal deficiency. Of particular interest, various investigators have shown that PKP2 deficiency leads to an increase in oxidative stress, ^5,6^ changes in the integrity and structure of the nuclear envelope^5^ and increased abundance of the immunoreactive signal for □H2AX, a marker of DNA damage repair (DDR). ^5^ These results, together with observations indicating an inflammatory component to the desmosomal arrhythmogenic cardiomyopathy phenotype^8–12^ are consistent with features that can also be observed in cases of cellular aging and senescence. ^13,14^ Yet, a focused study to test for the presence of senescence-associated structural and molecular features in desmosomal arrhythmogenic cardiomyopathies remains lacking. Identifying areas of convergence with other conditions related to senescence and premature aging (e.g., neurodegernative diseases; cancer) may guide novel approaches to therapy for PKP2-ARVC.

For the present study, we utilized a previously characterized murine model of PKP2 deficiency, namely, a tamoxifen (TAM)-activated, cardiomyocyte-specific PKP2 knockout (PKP2fl/fl/□MHC-CreERT2wt/+; referred to as PKP2cKO). ^15^ Following TAM injection, these mice develop, in a compressed timeline, the different phases of PKP2-ARVC: from a seemingly healthy heart to an arrhythmogenic cardiomyopathy of right ventricular predominance 21 days post injection (dpi) of TAM, followed by LV involvement (at 28 dpi), progressing toward end-stage bi-ventricular failure and death, with a 100% lethality at 50-60 dpi. ^15^ The reproducible sequence of events allows us to obtain molecular snapshots at specific timepoints after TAM, and relate those to the clinical phenotype.

An important feature of our cardiomyocyte-centric animal model is that it allows us to define the role of cardiomyocytes as a root source for the overall cardiac phenotype in the setting of PKP2 deficiency. Here, we exploited this characteristic to demonstrate morphological, biochemical, transcriptional and epigenetic features of premature aging and senescence in non-myocyte cells dissociated from PKP2cKO hearts either 21 or 28 days post-TAM injection. Interestingly, some canonical features of senescence were not observed in the cardiomyocytes themselves, though signs of premature aging were indeed present. Overall, our data strongly place the cellular and molecular phenotype of PKP2-deficient cardiac cells within the framework of a prematurely aging, pro-inflammatory, senescent heart. The implications of these findings in the context of potential therapeutic approaches are discussed.

## METHODS

Animal experiments were conducted in 3–6-month-old mice expressing a cardiomyocyte-specific deletion of Pkp2 (PKP2cKO). All experiments in PKP2-deficient mice were controlled with TAM-injected fl/fl, Cre-negative sex- and age-matched mice from the same colony. Of note, the αMHC-Cre-ER(T2) used for these experiments (starting with^15, 16^) is not the commercially-available *Mer-Cre-Mer* (B6.FVB(129)-*A1cf^Tg(Myh6-cre/Esr1*)1Jmk^*/J). Our Cre was generated by Takefuji et al^17^ and shown to not have cardiac effects of its own at the TAM doses that we use and within the time window that we study (up to 28 days after TAM). In fact, it is worth noting that a TAM-treated heterozygous PKP2cKO mice (i.e., αMHC-Cre-ER(T2)/*Pkp2* wt/fl) shows no phenotype (See Supplemental Figure I of^18^). Both female and male mice were included since previous observations have shown no sex-specific differences in the phenotype (see Figure S16 of^5^).

Procedures complied with NIH guidelines and were approved by the NYU IACUC (IA16-01021). All procedures are detailed in the Supplemental Material. Briefly, ventricular myocytes and non-myocytes were isolated by enzymatic dissociation using Langendorff perfusion. Non-myocytes were separated from myocytes by centrifugation and then either fixed in PFA for immunostaining or cultured for 24 hours. Images of non-myocytes were obtained by confocal microscopy. Conditioned medium was collected for cytokine profiling using the Proteome Profiler Mouse Cytokine Array. CpG island DNA methylation profiling was carried out by the Clock Foundation, using the Hovarth 320K Mammalian Methylation array. ^19^ Ventricular myocytes were imaged by conventional confocal microsopy or by a combination of expansion and Structured Illumination Microscopy (Ex-SIM²) to determine their molecular architecture at ∼40 nm optical resolution (primary antibodies listed in **Table S1**). Multiplex Opal 6-plex staining was performed on formalin-fixed paraffin-embedded heart tissue to identify the location of non-myocytes within a tissue section (antibodies listed in **Table S2**). Spatial transcriptomics was then performed on the same blocks using the Nanostring GeoMx platform; myocyte-rich and non-myocyte regions from the right and left ventricular free walls were segmented and analyzed.

### Statistical Analysis

Numerical results are reported as mean ± standard deviation (SD). Data pooled from individual animals were evaluated using the workflow described by Sikkel and collegues^20^ to assess the validity of the assumption of independence in the presence of potential data clustering. When clustering was detected, hierarchical analysis was performed according to the methods detailed in the supplementary material of^20^. All datasets were tested for normality using the Shapiro-Wilk test, and statistical significance was determined using either parametric or non-parametric tests, as appropriate. The specific tests used are indicated in the figure legends. Data analysis was conducted using RStudio and GraphPad Prism versions 9.0 and 10.0.

## RESULTS

### Senescence and premature aging in non-myocyte cells from PKP2cKO hearts

In the first part of our study, we examined whether the loss of PKP2 exclusively in myocytes was sufficient to induce a senescence phenotype in non-myocyte, cardiac resident cells. These cells were studied collectively in culture, after separation from the myocytes. Confocal microscopy showed that in PKP2cKO hearts, there was an abundance of non-myocyte cells with larger nuclei and dense heterochromatin aggregates, consistent with the description of senescence associated heterochromatin foci (SAHFs; **Figure 1A**). These heterochromatin densities were found in cells of various morphologies, including (but not limited to) those immuno-reactive to α-smooth muscle actin (**Figure S1**) and also those immuno-reactive to vimentin (**Figure S2**), a protein found in several non-myocyte cells including myofibrobasts and activated macrophages. Plots correlating the area of the nuclei to the number of heterochromatin foci per cell are shown in **Figure 1B-D**. In control hearts, approximatey 95% of the cells studied showed nuclei smaller than 75 µm² (**Figure 1B**) and therefore that value was taken as a parameter to compare against data collected from PKP2cKO 21 dpi and 28 dpi hearts (**Figures 1C** and **1D**, respectively). The analysis showed an abundance of cells with larger nuclei in PKP2cKO hearts. Importantly, SAHFs were abundant not only in cells with larger nuclei, but also in cells with nuclear size similar to control. Indeed, for nuclei smaller than 75 µm², the number of SAHFs significantly increased from an average of 2.48 ± 1.80 (n = 567 nuclei) to 3.43 ± 2.53 (n = 645) in PKP2cKO 21 dpi cells, and to 3.61 ± 2.49 (n = 1346) in PKP2cKO 28 dpi cells (**Figure 1E**). DAPI-stained cardiac tissue sections collected from PKP2cKO hearts revealed cells with abundant dense heterochromatin foci, thus supporting the notion that the observed densities in the isolated cells were not an artifact of the dissociation procedure (**Figure S3**).

**Figure 1.**
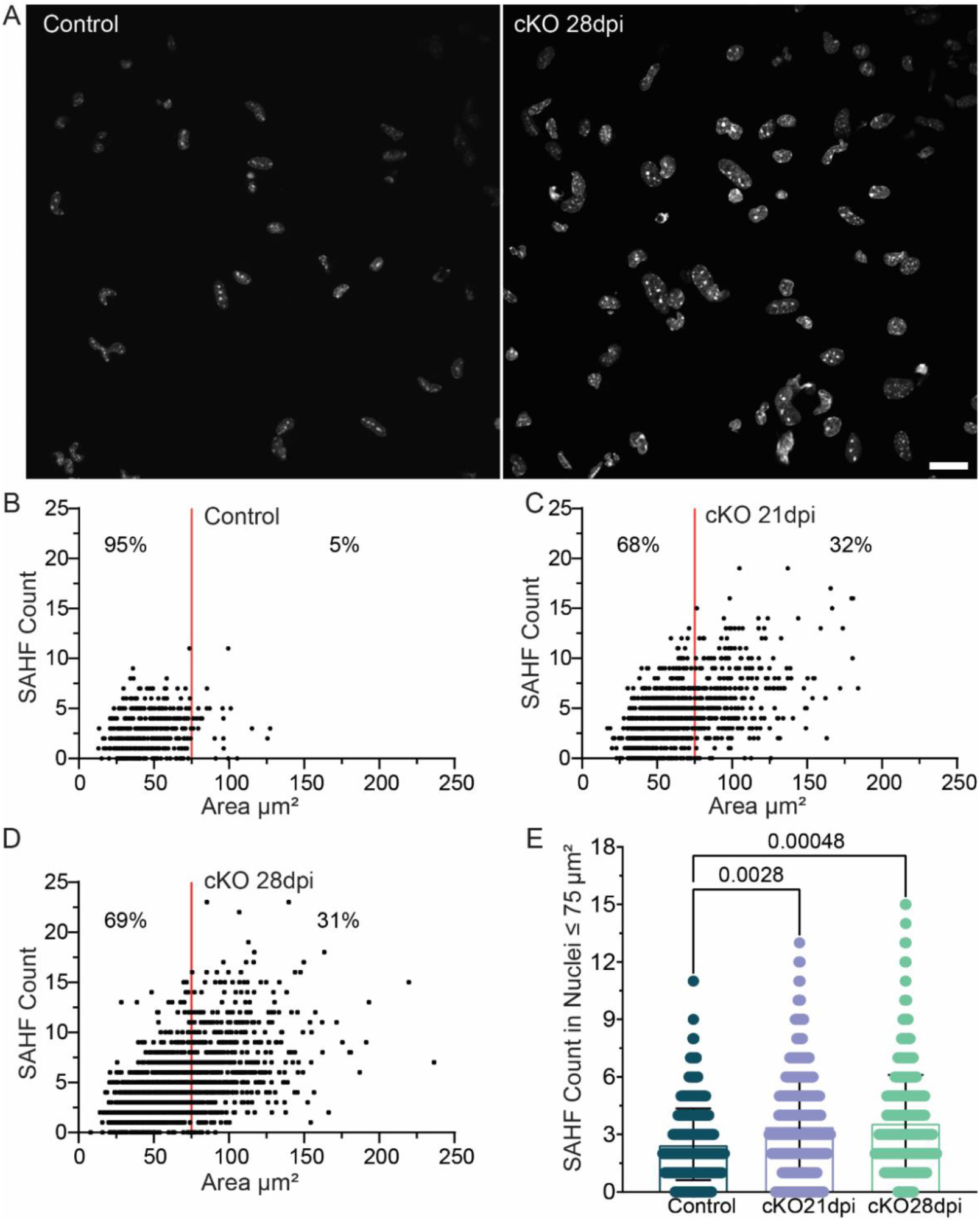
Heterochromatin densities in isolated PKP2cKO non-myocytes. (**A**) Nuclei from control (left) and PKP2cKO 28dpi (right) non-myocytes stained for DAPI (gray). Scale bar: 10 µm. (**B–D**) SAHFs as a function of nuclear area in control (**B**), PKP2cKO 21dpi (**C**), and PKP2cKO 28dpi (**D**). Red vertical line at 75 µm² marks limit containing 95% of data in control. Compiled data in (**E**). Data shown as mean ± SD; each point represents one nucleus. Minimal clustering was detected (ICC =1.8%) as per Sikkel et al. with a significantly better fit using multilevel modeling (p < 0.0001). ^20^ Thus, analysis accounted for clustering, followed by Bonferroni’s multiple comparisons test. Number of datapoints/mice: 567/5; 645/4; 1346/5 (control, PKP2cKO 21 dpi; PKP2cKO 28 dpi respectively).

Separate staining of isolated non-myocytes showed abundance of cells positive for p21, a protein commonly used as a marker of cell division arrest and senescence^21^ (**Figure S4**). Furthermore, we probed the media of PKP2cKO non-myocyte 24-hour cultures for the presence of cytokines previously associated with the senescence-associated secretory phenotyope (SASP). A total of 21 out of 40 spots in a commercially-available cytokine array yielded a signal; of them, 8 were significantly more abundant and 2 significantly less abundant in the media of cultures containing PKP2cKO non-myocytes collected at 21 dpi (**Figure 2**; SASP-related, as previously defined, ^22^ are underlined in red). Other cytokines were detected in samples from PKP2cKO cell cultures but not in controls. These included IL-1***α***, IL-16 and IL-17, which are part of the canonical SASP, or their downstream effects (**Figure S5**). A complete list is provided in **Table S3**. Cytokine abundance was either maintained or in some cases, reduced, when quantified at 28 dpi. Of note, as opposed to non-myocytes, cardiac myocytes cannot be cultured for the same time (24 hours) while preserving their morphology and function. As such, analysis of SASPs was conducted only at the transcript level (see below).

**Figure 2:**
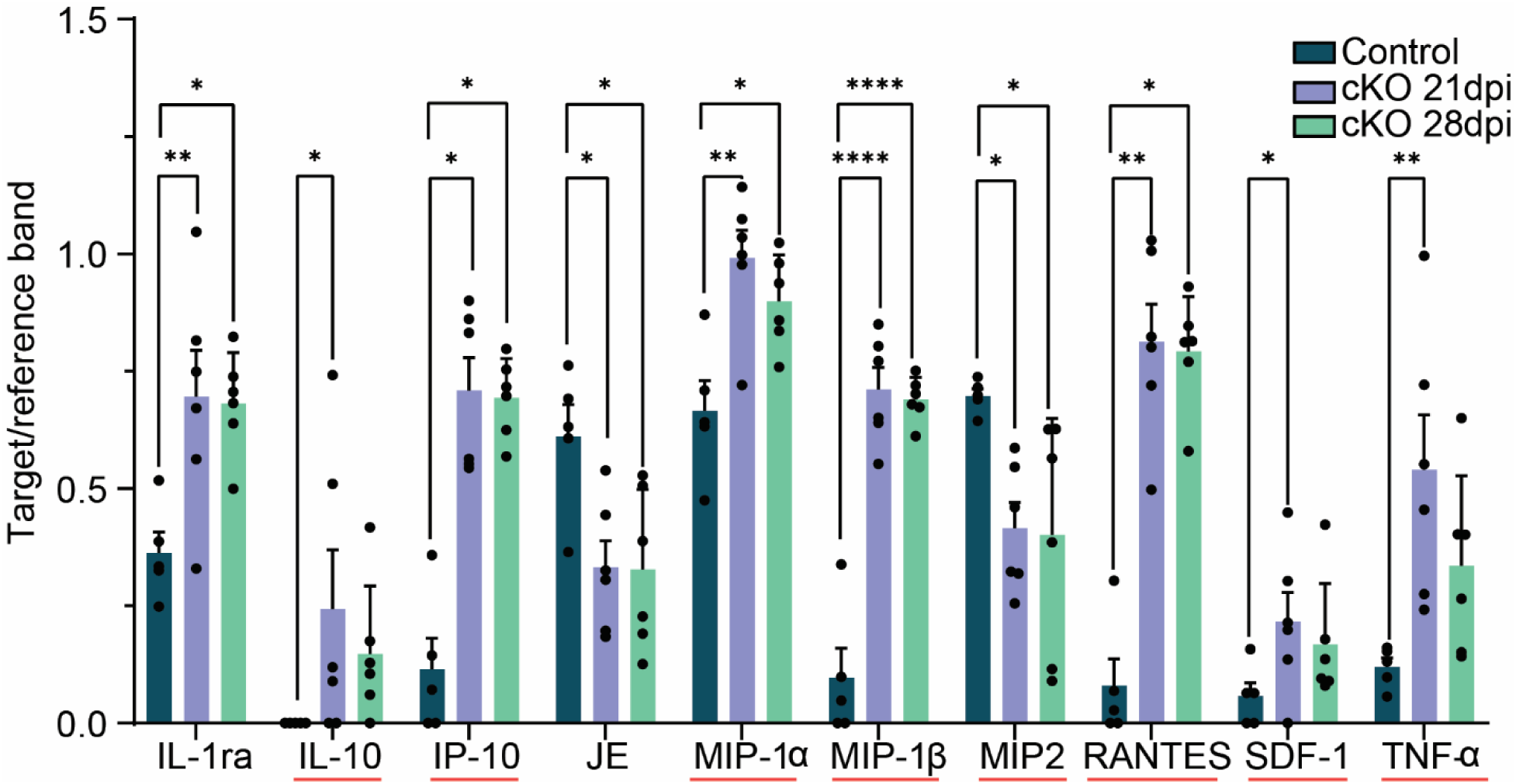
Cytokines in media of PKP2cKO non-myocytes in 24hr culture. Densitometry measurements of target molecules relative to reference band. Test molecules in abscissae; red lines mark canonical SASPs, as per Saul et al. ^22^ Data as mean ± SD. One point per mouse (n=6 per condition). Only cytokines appearing in control sample and showing statistical significance in test groups are shown. Normality assessed by Shapiro–Wilk test. Significance by ANOVA or by a non-parametric test, as appropriate (see **Table S3**).

Overall, the data presented above suggested that loss of PKP2 in myocytes can lead to a senescent phenotype in non-myocytes. Next, we investigated if, in addition, the PKP2 deficiency would trigger an increase in biological age in the non-myocytes. Non-myocyte cells were collected from PKP2-deficient hearts 28 days post-TAM, and their DNA methylation patterns were analyzed. ^19^ Four different, previously validated epigenetic or DNA methylation clocks were implemented including two different Pan-Universal Clocks (**Figure 3A and 3B**), the DNAmAge Heart Clock (**Figure 3C**) and the DNAmAge Fibroblast Clock (**Figure 3D**). ^23,24^ Consistently, PKP2cKO samples showed DNA methylation profiles matching those of older standardized samples, and significantly “older” than control. Analysis against additional clocks yielded similar results (**Figure S6**). It is important to note that there is no clock specifically outfitted for murine non-cardiomyocytes analyzed as a heterogeneous population. As such, the relevant parameter is not the absolute value of predicted age for the control cells but the differential between the age predicted by the clock for the control sample, and that determined for the PKP2cKO cells.

**Figure 3:**
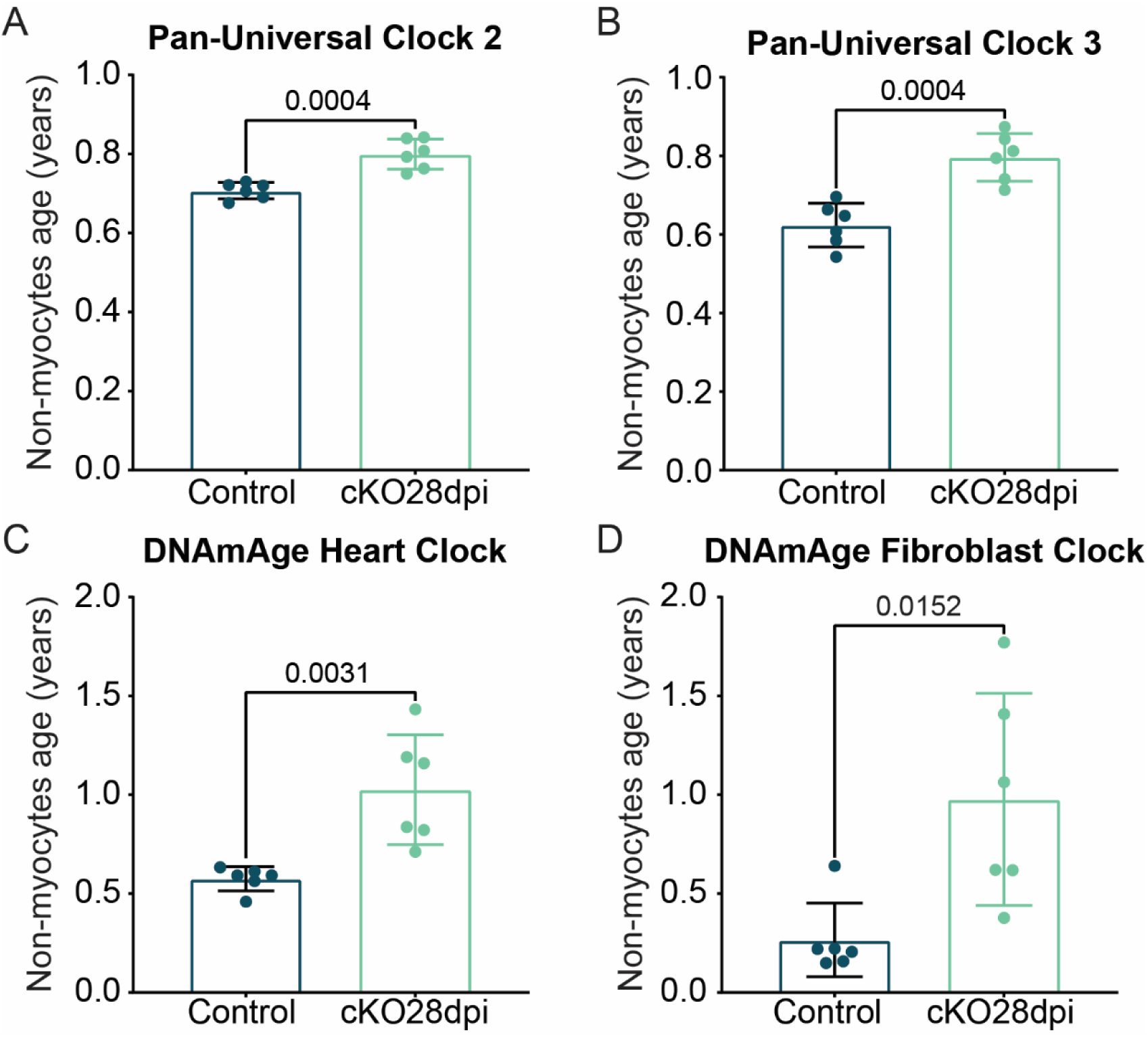
Predicted biological age of non-myocytes. DNA extracted from control and PKP2cKO 28 dpi non-myocyte samples, enzymatically isolated and immediately snap-frozen, was analyzed for their CpG methylation profiles (epigenetic clocks). (**A–B**): Pan-mammalian (Pan-Universal) clocks 2 and 3 (predict biological age across all mammalian species and all tissues^24^). (**C-D**) Mouse-specific clocks: (**C**) heart-specific and (**D**) fibroblast-specific. ^23^ Normality by Shapiro– Wilk test. Statistical significance by t-test for panels A-C and by Mann–Whitney test for panel D. Data shown as mean ± SD. One point represents one mouse (n = 6).

### Evidence of senescence in PKP2cKO tissue samples by spatial transcriptomics

The data described above was acquired from cells after isolation. We then asked whether the senescent phenotype would be present in cells in situ (i.e., not subjected to a dissociation protocol). As a first step, we examined the distribution of myocytes and non-myocytes in tissue sections of PKP2-deficient hearts. Multiplex immunochemistry of a PKP2cKO 28dpi heart showed that areas rich in myocytes also included cells positive for F4/80, Alpha-SMA, Ly6G, CD3, and CD11c (identifying respectively: macrophages, smooth muscle cells and activated fibroblasts, neutrophils, T lymphocytes, and dendritic cells; **Figure 4**). In contrast, in control hearts we only identified isolated macrophages and blood vessels positive for α-SMA (Figure S7). The antibodies used for probing were chosen given previous studies noting the importance of macrophages, activated fibroblasts and immune cells in the phenotype of desmosomal arrhythmogenic cardiomyopathies, ^10^ and taking into account the limited number of cell types that can be probed on the same sample. Overall, non-myocyte cells were found in niches, as well as intermingled in myocyte-rich areas (**Figure S8**). A separate example, obtained at 21dpi, is shown in **Figure S9A** and **S9B**. Considering that these areas may be the ones most influenced by the loss of PKP2 in myocytes given their physical proximity, we utilized the GeoMx platform to examine their transcriptomes, utilizing sections immediately subjacent to the ones used for the immunochemistry. Volcano plots are shown in **Figure 5A-D** (21 dpi, LV and RV in panels A and B, respectively; 28 dpi, LV and RV in panels C and D, respectively). Our transcriptomics data were cross-referenced against the SAUL_SEN_MAYO signature database for senescence and SASP. ^22^ SASP-related transcripts^22^ are marked in green in **Figure 5**. Other senescence-associated transcripts (though not specifically cataloged as SASP) are marked in the volcano plots in **Figure S10**. Our results show that a total of 47 out of 77 transcripts identified in the SAUL database as SASP-related were over-represented, and 2 were under-represented (using log2Fc=0.2 and P-adj=0.05 as criteria) in LV and/or RV samples collected from PKP2cKO hearts at 21 dpi (the complete datset is provided in **Table S4**). Interestingly, the number decreased to 21 transcripts for samples collected at 28 dpi (19 over-represented and 2 under-represented), suggesting not only a physical but also a temporal variation in the microenvionment as a function of disease progression.

**Figure 4.**
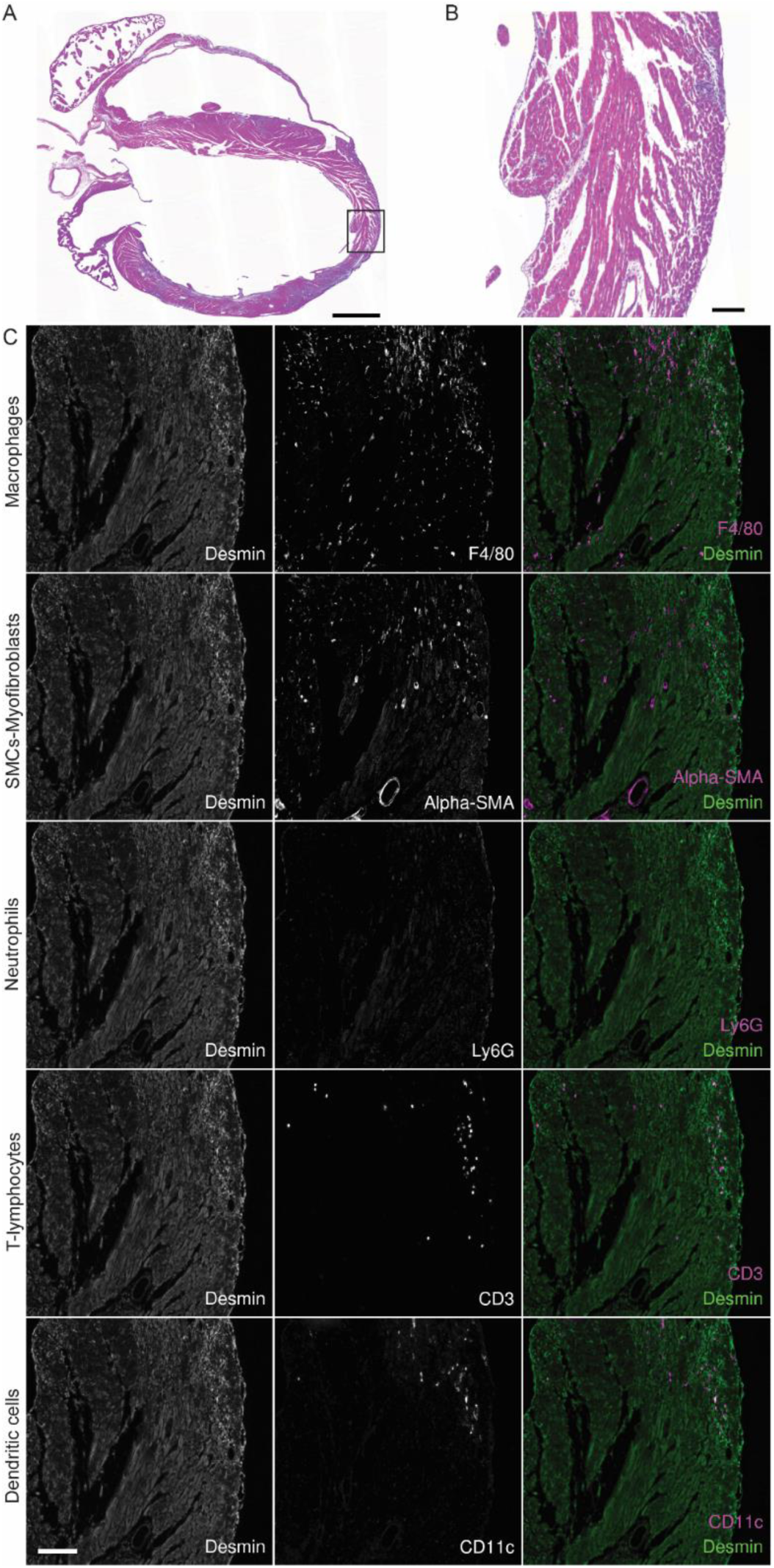
Multiplex immunohistochemistry of a PKP2cKO 28 dpi heart. Hematoxylin and eosin (H&E) staining is shown in panel **A**; small square marks the area shown in **B** at higher magnification. Panel (**C**) shows immunohistochemistry of the immediately subjacent section, focusing on the area in B, using an Opal 6plx staining to mark macrophages (F4/80), myofibroblasts (alpha-SMA), neutrophils (Ly6G), T-lymphocytes (CD3) and dendritic cells (CD11c). Corresponding images are shown in the middle panels. Section was also stained for desmin (left panels) to highlight myocytes (desmin-positive and yet negative for all other markers). Merged images shown in the right. Scale bars: 1 mm (A); 100 µm (B, C).

**Figure 5.**
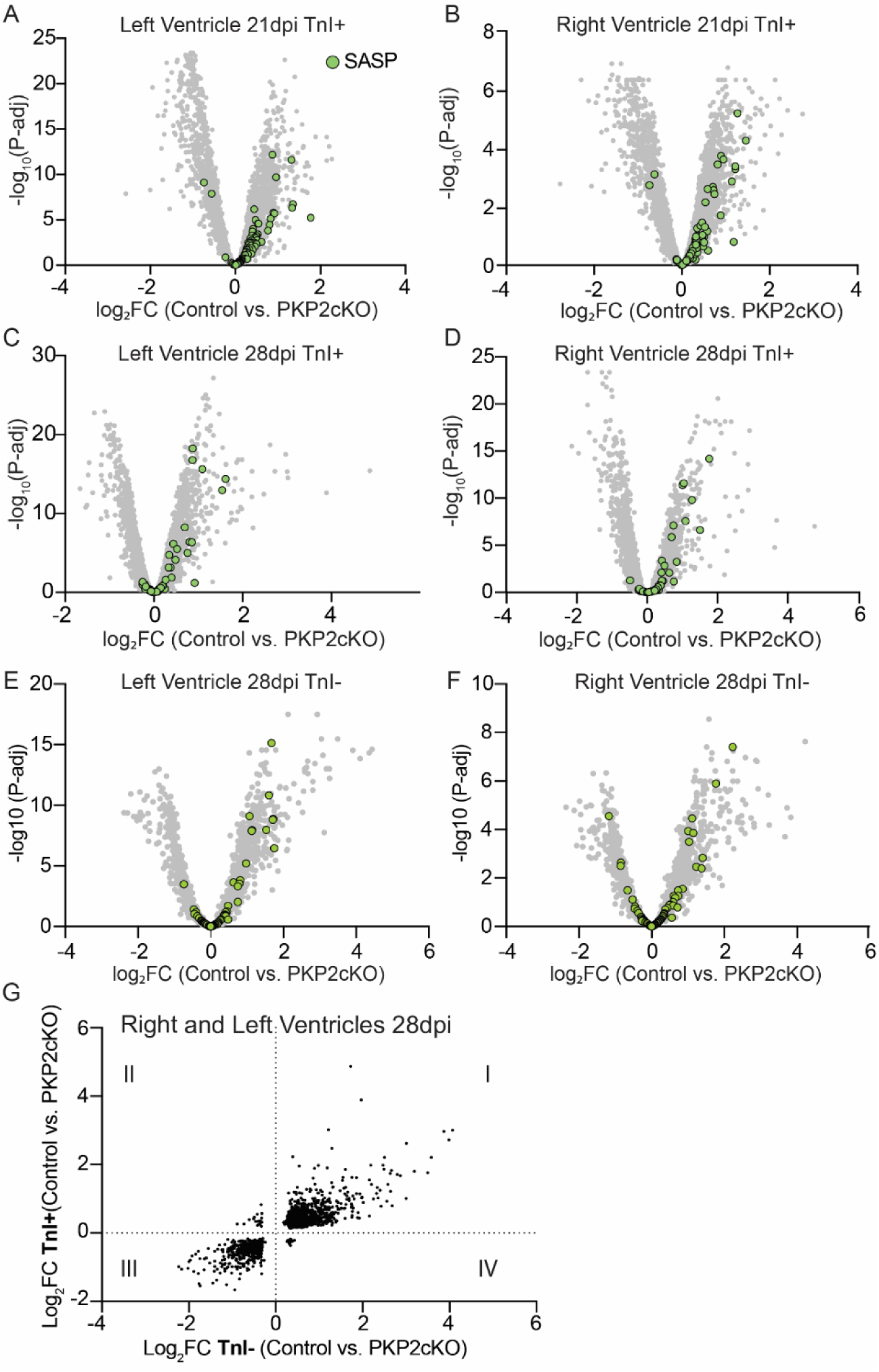
Region-specific differential transcriptomes. Each Volcano plot correlates magnitude of the differential (Log2Fc) against false discovery rate (P-adj) per gene. Gray dots: differentially expressed transcripts. Green: Canonical SASP-related transcripts, as per Saul et al. ^22^. TnI+ or TnI-indicate regions abundantly marked, or prominently void, of a Troponin-I signal, respectively. Samples collected from left and right ventricle of PKP2cKO hearts 21 (panels **A-B**) or 28 dpi (panels **C-F**) analyzed by the GeoMx platform. Total numbers of mice: 12 for control (6 parallel to 21dpi and 6 parallel to 28dpi), 6 for PKP2cKO 21 dpi and 6 for PKP2cKO 28 dpi. Data collected from 7 to 12 ROIs in the right or left ventricular free wall. **G**: log2Fc collected from TnI- (abscissae) or TnI+ (ordinates) at 28 dpi. Each data point represents one transcript. Quadrants divided by the Cartesian map origin (0,0) and marked by roman numerals.

The GeoMx platform also allowed us to examine the transcriptomes in areas where myocytes were not detectable (i.e., negative for TnI staining). **Figure 5E** and **5F** show the volcano plots of the transcriptomes acquired from TnI- regions at 28 dpi (the limited size of TnI-areas at 21 dpi precluded us from proper transcriptomics analysis in that specific group, the complete datset is provided in **Table S5**). The data show SASP-related transcripts also in this population, though in lower numbers than in the 21 dpi TnI+ groups. Yet, when analyzed as a whole, there were strong similarities between the transcriptomes of TnI+ and of TnI- regions. This is emphasized by the plot presented in **Figure 5G**. Each data point corresponds to one gene and only genes with an FDR <0.05 are shown. The log_2_FC values for the transcriptomes obtained from TnI+ regions at 28 dpi are presented in the abscisa and those obtained from TnI- regions at the same time point are presented in the ordinate. The Cartesian plot is heavily populated in the first and third quadrants, indicating equal directionality of change for both cardiac regions (the entire list of genes present in each quadrant is reported in **Table S6**).

These results strongly support the notion that loss of PKP2 in cardiomyocytes is a sufficient condition to trigger a senescent phenotype in regions proximal, as well as distal to the myocytes, both populated with non-myocyte cells. As a next step, we explored signs of senescence or premature aging in the cardiomyocytes themselves.

### Cardiomyocytes exhibit signs of premature aging

Analysis of enzymatically dissociated cardiomyocytes did not show abundant expression of p21 or presence of SAHFs, neither in control nor in PKP2cKO cells (**Figure S11A-D).** Moreover, cardiomyocytes did not manifest DNA methylation signatures consistent with an elevated biological age compared to controls (**Figure S11E-H**). Yet, previous reports have shown other signs of premature aging in PKP2cKO cardiomyocytes including excess reactive oxygen species (ROS), abundant expression of H2AX (a marker of DNA damage response/repair, DDR), increased nuclear envelope permeability, and altered nuclear size and morphology. ^5^ To complete the structural profile of cardioyocytes from the perspective of premature aging, we explored additional specific features: chromatin distribution, and nucleolar morphology and function.

### Heterochromatin redistribution in PKP2cKO myocytes

Previous studies have shown that the spatial organization of heterochromatin is disrupted in aging myocytes. ^25^ In this study we utilized a combination of expansion microscopy and structured illumination (Ex-SIM^2^) ^26,27^ to investigate, at sub-diffracion limit resolution and in three dimensions, the chromatin architecture of adult ventricular myocytes. Details of the methodology are presented in **Figure S12** and **Videos S1**, **S2** and **S3**. Single optical slices through the midsection of nuclei from a control myocyte, or from a PKP2cKO myocyte, are shown in **Figure 6A**, DNA was labeled using SYTOX orange. Brighter staining corresponds to regions of more compacted chromatin, likely reflecting heterochromatin domains. The slices displayed were selected from the midsection of the nucleus, oriented parallel to its long axis, thus visualizing the lamin associated domain (LAD) as a ring of dense compacted chromatin around the nuclear contour (see diagram in **Figure 6B**). From these images, we quantified the SYTOX signal intensity at LADs relative to the total detected in the nucleus. We excluded the top and bottom slices, since in those cases the LAD runs parallel to the direction of the optical slice. We observed a reduction in SYTOX signal at LADs in PKP2cKO mice compared to controls at both 21 and 28 dpi, likely reflecting less compacted chromatin in these regions (**Figure 6C**).

**Figure 6.**
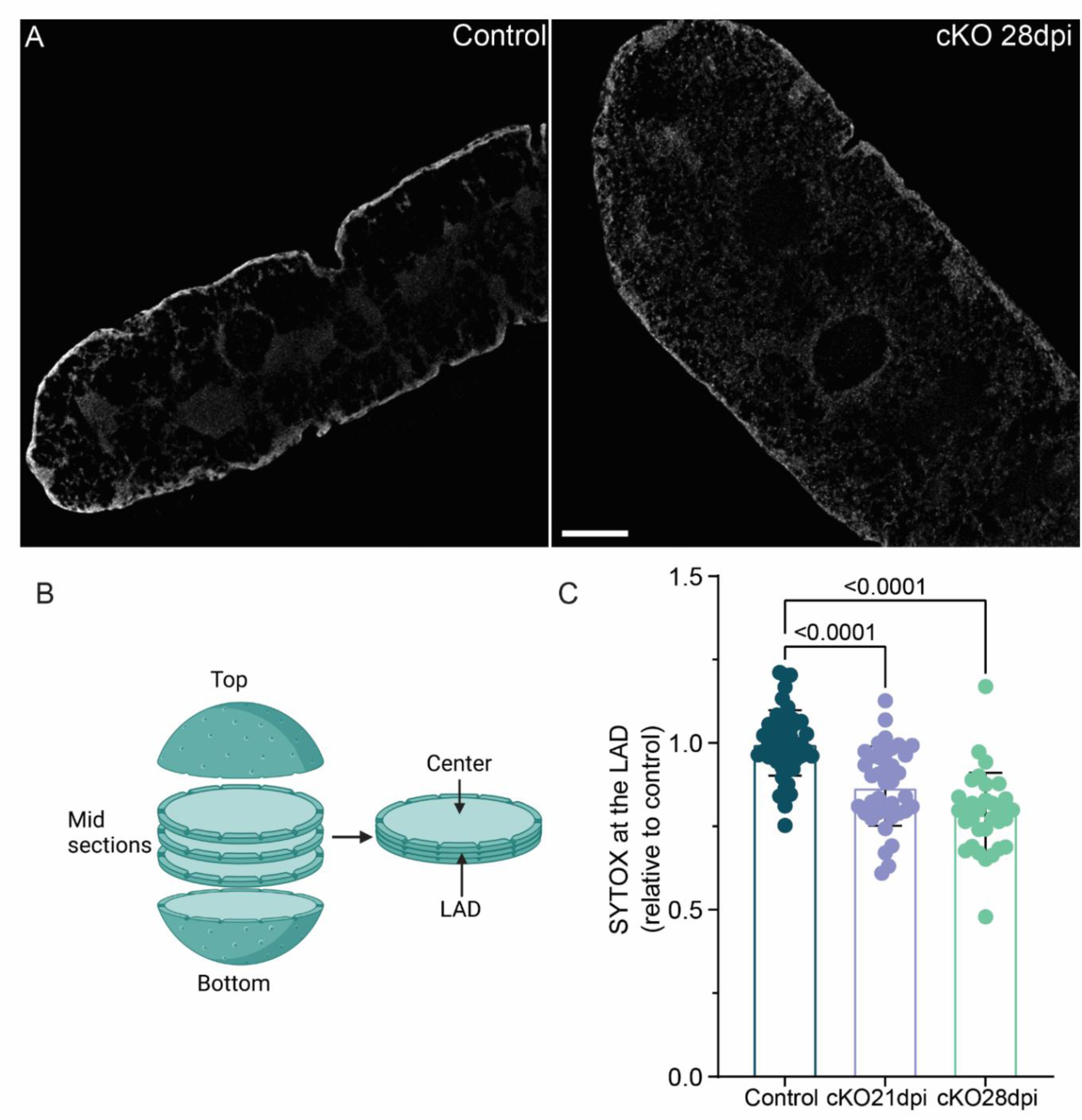
Chromatin remodeling in PKP2cKO ventricular myocytes. (**A**) Representative images of mid-section stacks (excluding top and bottom) of nuclei from control (left) and PKP2cKO mice 28dpi (right), stained with SYTOX orange. Cells subjected to expansion and imaged by SIM^2^ (ExSIM^2^; see Methods). Scale bar: 1.5 μm. (**B**) Diagram illustrating the planes selected for analysis. (**C**) Proportion of dense chromatin (by SYTOX) at the nuclear perimeter (lamin-associated domain; LAD) relative to total SYTOX-positive signal in control, PKP2cKO 21 dpi, and PKP2cKO 28 dpi. Data normalized to the average control value at each experiment. One point represents one cell; n/N=48/7, 38/6 and 32/4 for control, PKP2cKO 21 dpi and PKP2cKO 28 dpi respectively (n=number of cells; N= number of mice). Vertical thin lines indicate SD. Clustering was negligible (ICC < 5%, no superior multilevel fit). ^20^ Normality assessed by Shapiro–Wilk; significance by one-way ANOVA followed by Bonferroni for multiple comparisons.

### Nucleoli as a reporter of premature aging

Previous studies have found nucleoli morphology and nucleoli transcriptional activity to be reporters of premature cell aging. ^28–30^ We therefore directed our Ex-SIM^2^ system to characterize, at subdiffracion limit resolution, the anatomical features of nucleoli in PKP2cKO myocytes. **Figure 7A**, **Videos S4** and **S5** show examples. Dimensions were collected from the optical slice that revealed the largest nucleoli internal diameter, measured to the heterochromatin contour. Nucleoli were larger in PKP2cKO myocytes collected both at 21 dpi and at 28 dpi (**Figure 7B**; circularity measures in **Figure S13**). We also observed that in control myocytes fibrillarin organized in rosettes, harboring H2AX, but this organization was disrupted in the PKP2cKO cells (bottom panels in **Figure 7A**). Also of note, the abundance of H2AX trended toward an increase (**Figure 7C**), as it also occurs in cells undergoing nucleolar stress. ^29,31^ When observing at super-resolution (around 40-60 nm in our case), the signal of DAPI and that of H2AX did not overlap, as expected given that DNA damage repair would not be occurring at a site of condensed heterochromatin.

**Figure 7.**
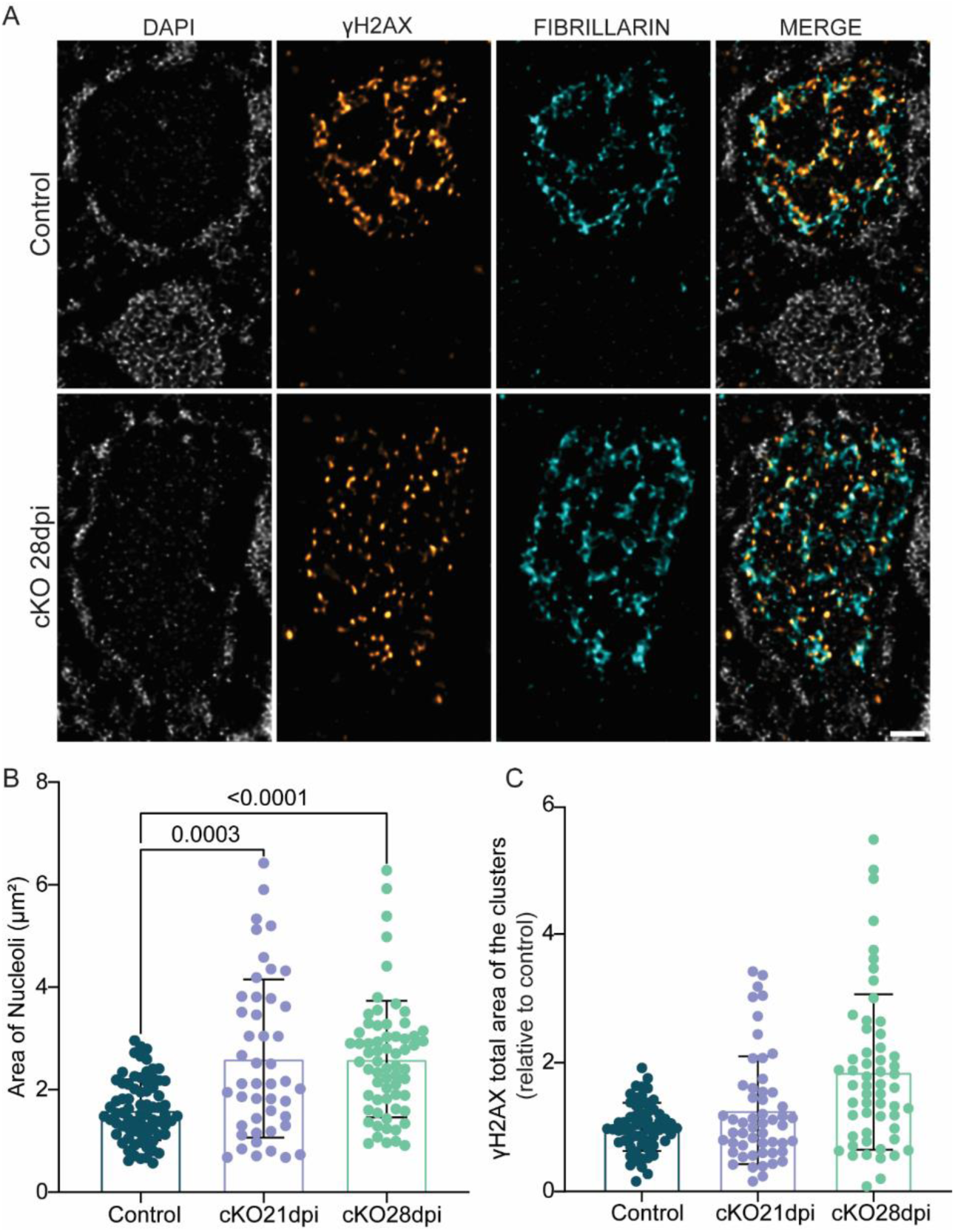
Nucleolar architecture and γH2AX abundance in PKP2cKO cardiomyocytes. (**A**) Nucleoli from control (upper panel) and PKP2cKO 28 dpi (lower panel). White: DAPI (left panels). Gold and Cyan: γH2AX and fibrillarin, respectively. Right: merged images. Scale bar 300 nm. See also **Videos S4/S5**. **B**: Nucleoli area (at optical stack midsection). One point per cell; number of cells/Number of mice=74/9, 46/5 and 64/7 for control, PKP2cKO 21 dpi and PKP2cKO 28 dpi respectively. Vertical lines mark SD. Clustering was negligible (ICC < 5%, no superior multilevel fit). ^20^ Normality assessed by Shapiro–Wilk test; significance by Kruskal–Wallis followed by Dunn’s multiple comparisons. (**C**) Area occupied by γH2AX within nucleoli. Data normalized relative to average control; mean ± SD. One point per cell. n/N=60/7, 53/6 and 56/5 for control, PKP2cKO 21 dpi and PKP2cKO 28 dpi respectively. Clustering analysis^20^ revealed intraclass correlation (ICC = 57.5%) and significantly better fit by group-level model (p < 0.0001); statistical significance assessed accounting for clustering. ^20^ No differences across conditions but a trend toward increased DDR area in PKP2cKO 28dpi (p=0.07 Control vs PKPcKO28dpi).

Control experiments using Neocarzinostatin (NCS), a potent DNA damage inducer, ^32^ showed the expected increase in γH2AX in the nuclei. When measured separately, the increase was apparent in the nucleoplasm (excluding the nucleoli), but not in the area within the contour of the nucleoli (see Figure S14A-C; nucleoli highlighted in yellow; see also Perez-Hernandez et al., ^5^). Separately, we examined the abundance of upstream binding factor (UBF) as a marker of ribosomal DNA transcription in the fibrillar center. ^33^ The data showed increased abundance of UBF in nucleoli from PKP2cKO myocytes when compared to control. Individual examples are shown in **Figure 8A** (notice the distrupted organization of fibrillarin rosette, as also observed in the previous example) and the quantitative analysis in **Figure 8B and 8C**. Overall, we found nucleoli of PKP2cKO myocytes to be larger, of disrupted architecture and increased transcriptional activity, when compared to control. Based on these results, we targeted our transcriptomics analysis to evaluate the magnitude of ribosomal RNA in relation to PKP2 deficiency. The data showed that 30 out of 48 ribosomal transcripts (as per the Ribosomal Protein Gene Database) ^34^ were over-represented in the PKP2cKO transcriptomes, both at 21 and at 28 dpi (**Figure 8D, Table S7**). Similar results were obtained when RV samples were analyzed (**Figure S15** and **Table S7**). These results were in agreement with previous studies on nucleoli transcriptional activity in premature aging cells. ^28,29^

**Figure 8.**
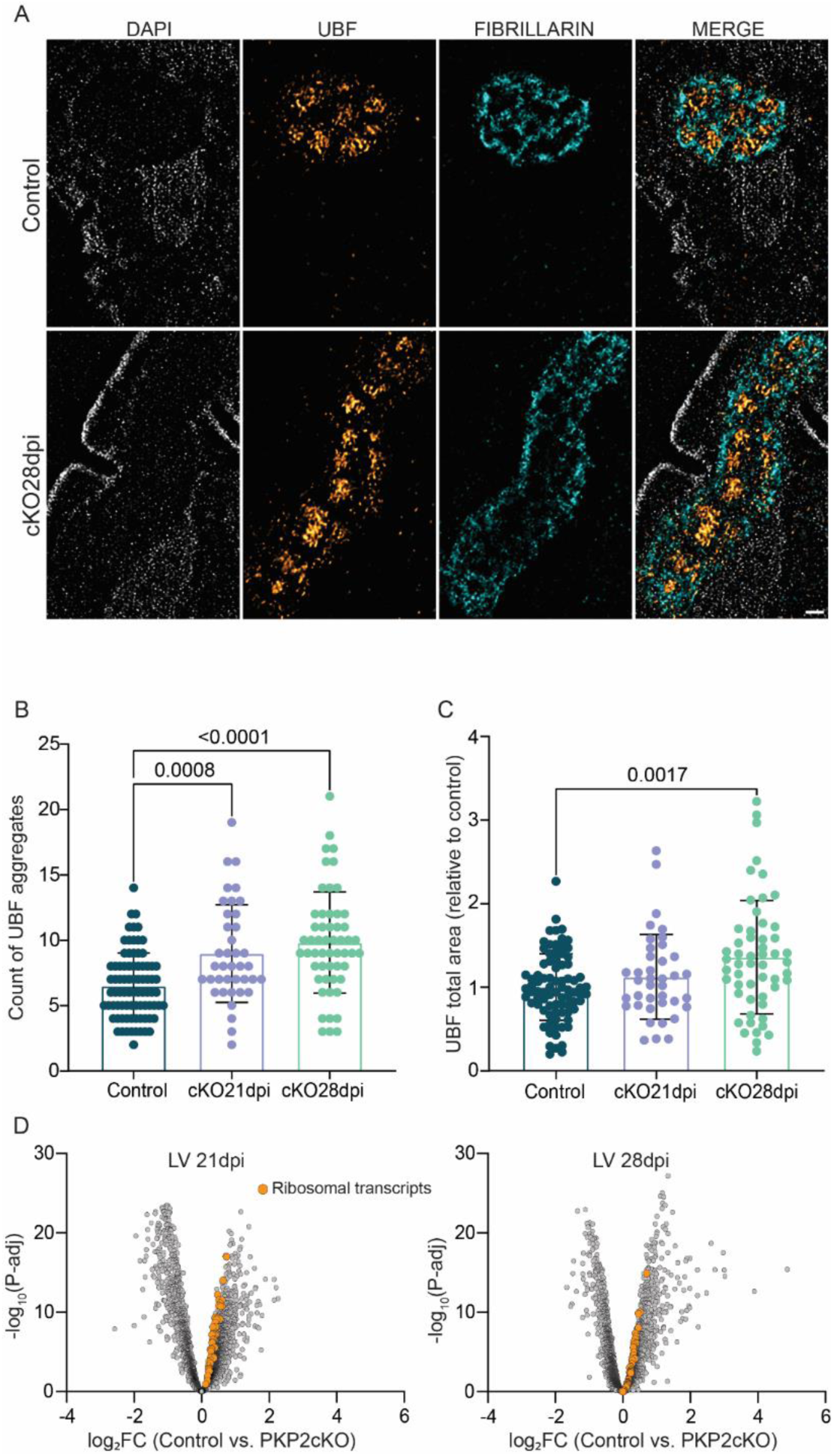
Transcriptional activity in nucleoli of PKP2cKO cardiomyocytes. (**A**) Nucleoli from control (upper panel) and PKP2cKO 28 dpi (lower panel) stained for DAPI (white), UBF (gold), and fibrillarin (cyan). Merged image in the right. Scale bar 300 nm. (**B**) Number of UBF aggregates in nucleoli. (**C**) Area occupied by UBF signal, normalized to average control value for each experiment. Data shown as mean ± SD. One point per cell. number of cells/Number of mice: 82/8, 39/4 and 54/6 for control, PKP2cKO 21 dpi and PKP2cKO 28 dpi respectively. Negligible clustering (ICC < 5%, no superior multilevel fit). ^20^ Normality of distribution by Shapiro–Wilk; significance by Kruskal–Wallis followed by Dunn’s multiple comparisons. (**D**) Volcano plots for LV 21 dpi and LV 28 TnI+ differential transcriptomes (same as in Figure 5A and 5C) marking ribosomal transcripts (as cross-referenced to the Ribosomal Protein Gene Database) ^34^ in orange.

### Correlative studies supporting the notion of an aging phenotype in PKP2cKO hearts

For further examination of the relation between cardiac aging and PKP2 deficiency, we cross-referenced the proteome of the PKP2cKO mouse at 21 dpi^5^ with the proteomics analysis of the heart of aging mice, the latter as defined by Takasugi and colleagues from the comparison of the cardiac proteome of 30-month old mice against that of 6 month-old mice. ^35^ The analysis showed that over 200 dysregulated proteins consequent to loss of PKP2 expression are also significantly different, at a false discovery rate (expressed as P-adj) of <0.005, in the aging heart (listed in **Table S8**). The number of convergent results increased to 488 proteins when the threshold was lowered to an FDR of <0.05 (**Table S8**). The Cartesian plot presented in **Figure 9A** shows the close correlation between the two datasets for data with FDR<0.005, strongly indicating that the PKP2cKO mice, which have a chronological age of ∼4-5 months, have a differential proteome expected for a mouse over 5-6 times their chronological age.

**Figure 9.**
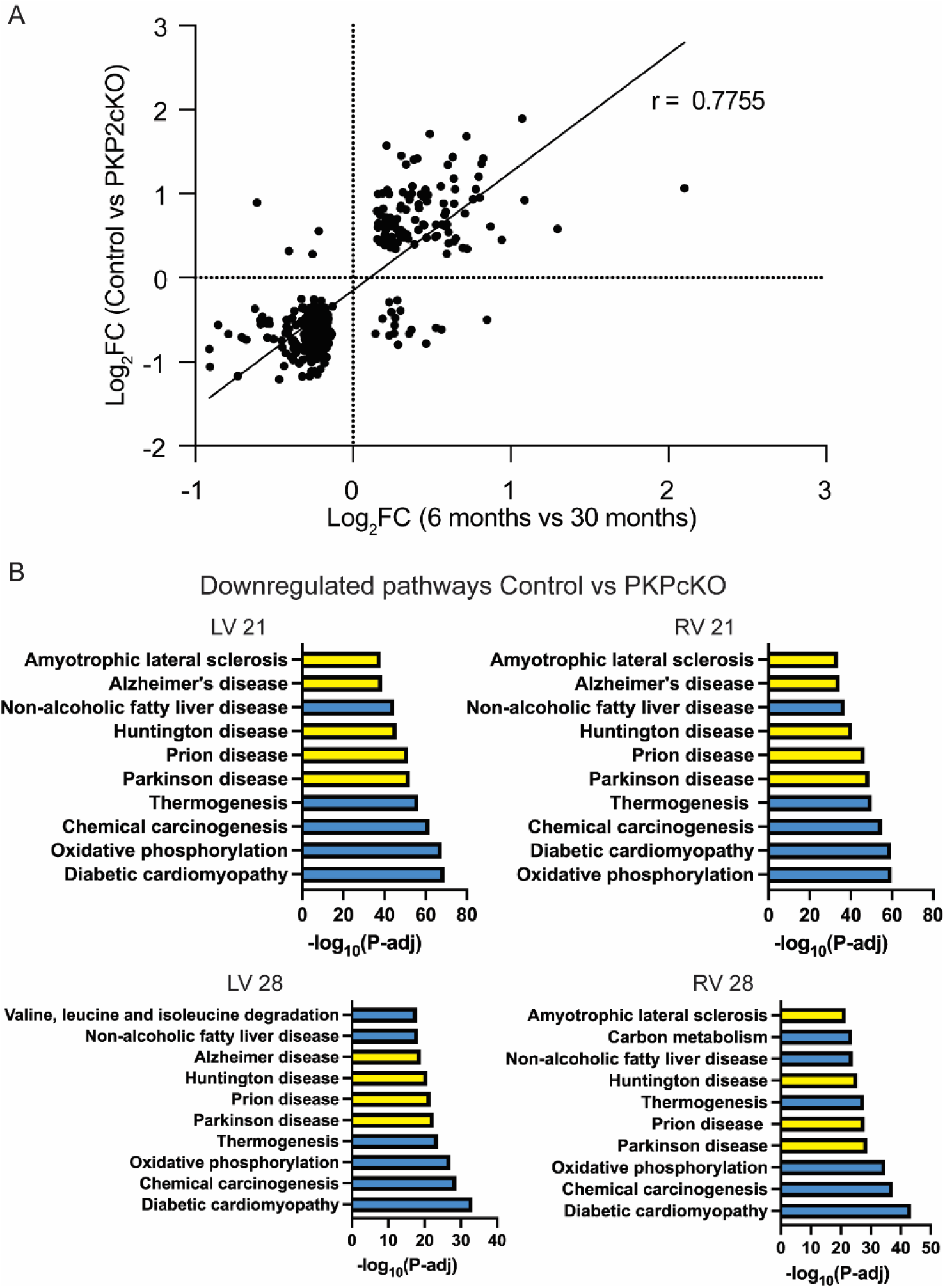
Cross-referenced analysis of omics data from PKP2cKO hearts. (**A**) Correlation between differential protein expression in Control vs. PKP2cKO 21 dpi hearts (ordinates; data from^5^) and proteome of 6- vs. 30-month-old mouse hearts (abscissae; data from Takasugi et al^35^). Each point marks one protein. Crossing point of dotted lines marks origin of Cartesian plot (0,0). Only proteins with adjusted p-value < 0.0005 are included. Pearson correlation coefficient r = 0.7755. (**B**) Kyoto Encyclopedia of Genes and Genomes (KEGG) pathway analysis of downregulated transcripts in TnI+ regions from hearts collected from LV and RV at 21 dpi (top) and 28 dpi (bottom). Top 10 significant pathways, with neurodegenerative diseases highlighted in yellow. Thresholds of |log₂FC| > 0.5 and adjusted p < 0.005 were applied.

Pathway analysis of transcripts under-represented in PKP2cKO hearts (compared to control) revealed a significant enrichment in neurodegenerative disease pathways, particularly Alzheimer’s, Parkinson’s and amyotrophic lateral sclerosis (ALS). ^36^ This pattern was observed both in samples collected from TnI+ regions (LV and RV at 21 and 28 dpi; see **Figure 9B**) and in TnI– regions at 28 dpi (see **Figure S16**). For further confirmation, we cross-referenced our transcriptomic dataset against reference databases for neurodegenerative disorders, specifically Alzheimer’s disease, Parkinson’s disease and ALS. ^37^ A total of 176 transcripts matching the DISGENET database were dysregulated in the TNI+ LV 21 dpi group, and 73 of them were found only in that dataset (The specific genes are presented in **Table S9**). Whether this reflects generalizable cell biological processes being affected by unrelated pathological conditions, or a common pathway specific for degenerative diseases (neuro- or cardio-) will require further study.

## DISCUSSION

Age is a highly significant risk factor for death, regardless of cause. Previous studies show an acceleration of the aging trajectory consequent to chronic and acute diseases^46–48^ including degenerative conditions^49^. Acute stressors (e.g., major surgery, pregnancy, or COVID-19 infection) as well as chronic stress and disease (e.g., obesity and HIV infection), can increase biological age, measured at the molecular level, in various tissues and organs. ^38–42^ Senescence has been studied in the context of acquired heart disease^14^ and for inherited cardiomyopathies originating in the nuclear envelope. ^43,44^ To our knowledge, this is the first report linking a desmosomal arrhythmogenic cardiomyopathy to premature aging and to the induction of cellular senescence. Our results show that loss of PKP2 only in cardiomyocytes is sufficient condition for the induction of a pro-inflammatory senescent phenotype in non-myocyte cells, and for the premature aging of the heart.

Cellular senescence is defined as a biological state in which a cell permanently stops dividing but remains metabolically active. Features such as the presence of SAHFs, SASP and p21 are grouped under the broad description of senescence. ^45^ The accumulation of senescent cells in tissues is more prevalent in aged organisms and can itself promote cellular aging through the SASP. ^46^ Premature (biological) aging refers to the process when the molecular, phenotypic, and functional changes associated with aging appear earlier than expected for someone’s chronological age. While the latter purely reflects the time elapsed since birth, biological age is a dynamic reflection of the molecular and functional state of cells, tissues, and organs. ^13^ All organ-specific differences notwithstanding, our results lead us to conclude that premature biological aging and the pro-inflammatory senescent process are components of the molecular phenotype of PKP2-ARVC, partly sharing substrates with those observed in neurodegenerative diseases.

### Senescence in non-myocytes, inflammation and the molecular pathophysiology of PKP2-ARVC

Our data reveal molecular hallmarks of senescence and premature aging in non-myocytes, namely, the presence of SAHFs, a prominent SASP, expression of p21 and a significant increase in epigenetic age. Senescence is intimately related to the inflammatory process in a positive feedback loop where senescent features promote inflammation and the latter, in turn, accelearates senescence. This feedback loop is likely to contribute to the overall detriment of the cells and tissues, as it is observed in PKP2-ARVC. Our results therefore contribute to emerging evidence, following the seminal work of the Saffitz group^8–11^ indicating that the inflammatory and immune responses are key pathophysiological substrates of desmosomal cardiomyopathies. ^47^

We observed senescence in non-myocytes and yet, the loss of PKP2 occurred only in cardiomyocytes. The latter strongly indicates that myocytes can be primary triggers of the senescent response, likely by paracrine communication with non-myocytes in the vicinity. The identity of the “first responder” molecule or set of molecules released by the myocytes that initiate the global senescent/inflammatory process, remains unclear. Our efforts to obtain a spatial transcriptome that would encompass only myocytes were unsuccessful, as histological analysis revealed the intermingling of non-myocytes even in areas rich in TnI+ cells. Yet we speculate that the myocyte trigger is linked to the physical damage that occurs to the nuclear envelope and to the overall myocyte, following loss of intercellular adhesion. ^5^ It is also tempting to speculate that rather than exocytosed molecules, mitochondria, which are found in excess in PKP2-deficient myocytes and, occasionally, in their surrounding extracellular space (see Pérez-Hernández et al. ^5^) could act as triggers of the global inflammatory response, as it has been proposed for other systems. ^48,49^ Further studies will be necessary to test the latter hypothesis.

### Could senescence be a preamble to adiposis?

A characteristic of PKP2-ARVC is the abundance of adipocytes infiltrating the cardiac parenchyma; yet, the origin of the adipocytes remains unclear. A characteristic of senescence is the arrest of cell division. It is tempting to speculate that the arrest of cell division in activated fibroblasts becomes a permissive step for the transformation of the fibroblasts into adipocytes. Previous studies in cultured cells have suggested that epicardial-derived cells are the source of fibroblasts, and that they can transform into adipocytes, ^50–54^ though other studies have suggested that senescence impairs adipogenesis. ^55^ Murine models, including our own, have failed to show extensive adipocyte infiltrates, though the latter may be consequent to the short life-time of the mice vis a vis the time needed for extensive adiposis to manifest.

### SASP and arrhythmogenesis

The nature of the SASP is relevant not only from the perspective of the inflammatory response and its consequences on myocyte integrity but also, from the point of view of arrhythmogenesis. Indeed, multiplex immunohistochemistry showed that the distribution of specific non-myocyte cell types across the parenchyma is not homogeneous, thus suggesting that myocytes in one area may be exposed to different cytokines than myocytes in a neighboring region. As the impact of specific cytokines on cell electrophysiology differs, ^56^ we speculate that the heterogeneous distribution of non-myocytes can translate into electrical heterogeneity, yielding an overall arrhythmogenic substrate. Future studies will address this specific hypothesis.

### Cardiomyocyte structure and its relation to premature aging

#### Ex-SIM^2^

Changes in chromatin and nucleoli architecture are among the hallmarks of aging. ^13^ Proper characterization of those features required visualization by light microscopy at optical resolutions below the diffraction limit of light. In the present study we implemented a novel combination of expansion microscopy^26^ and structured illumination technology (Ex-SIM^2^). ^27^ With the expansion protocol we increased cell size by a factor of 6.4X. The persistence of regular z-disk separations, the regularity of the shape of the nuclei (in control cells) and the preservation of the overall cell geometry (including the desmin and alpha-actinin scaffolds) indicated that the expansion occurred in a proportional way. Expanded cells were then subjected to visualization and image reconstruction using the SIM^2^ technology. The calculated optical resolution with the latter method is ∼60 nm. ^57^ If divided by the expansion factor we could calculate that our resolution is near 10 nm. Even if overestimated, it is reasonable to consider that our optical resolution is better than 40 nm, the latter being the predicted range of influence of a molecule on its neighbors. ^58^ As such, our system allows a detailed analysis of molecular topology in the nucleus, and even within the nucleoli, which has an estimated diameter of ∼1.5 µm in a normal adult myocyte. ^59^

#### Chromatin remodeling at the LAD

We report a reorganization of chromatin at the expense of the lamin-associated domain in PKP2cKO cardiomyocytes. LAD reorganization has been reported as a feature of cell premature aging, ^25,60,61^ and it can result from an imbalance of the forces exerted on the nuclear envelope, as shown by Pérez-Hernández al. ^5^ PKP2-deficient myocytes may present a similar case. Indeed, previous studies show that in an adult cardiomyocyte, nuclear architecture is maintained by the balance of forces between desmin filaments pulling at the nuclear envelope, and dynamic microtubules pushing on the same structure^62^. We previously showed that loss of PKP2 leads to a separation of desmin from the nuclear envelope, thus impairing the nuclear membrane barrier and disrupting nuclear morphology. These changes in nuclear architecture are likely to cause physical stress on the heterochromatin located at the LAD, thus affecting its integrity and ultimately, its transcriptional activity. ^5^ The latter connects with the observation that mutations in lamin A/C, a protein of the nuclear envelope, can lead to rapid aging or to an arrhythmogenic cardiomyopathy with some features reminiscent of what it is observed in PKP2-ARVC. ^43^ Similar observations in models of TMEM43 deficiency connect nuclear envelope integrity with biological aging. ^44^

#### Nucleoli as reporters of premature aging

Ex-SIM^2^ observations revealed an increase in the size and a departure from circularity of the nucleoli, consistent with previous observations of nucleoli stress in a different heart disease. ^63^ We also report here an increase in the abundance of DDR signals, a disruption of the rosette structure formed by fibrillarin and an increase in a marker of active transcription (UBF), ^33^ which correlated with an increase in the abundance of ribosomal transcripts. All these elements combined were consistent with the notion of nucleolar stress, ^31^ a feature of premature aging. ^28,29^

### Cardiac premature aging and neurodegenerative diseases

KEGG pathways, and direct cross-reference analysis, indicate that a number of transcripts previously reported as dysregulated in neurodegenerative diseases are also dysregulated in the PKP2cKO hearts. This convergence was present when separately analyzing samples from the right or the left ventricle, and both at 21 and 28 days post-TAM. It is clear that neuro-degenerative diseases present hallmarks unique to neurons. Yet, it also seems clear that biological processes not unique to neurons are part of the molecular pathophysiology of PKP2-ARVC. These include, in myocytes: metabolic dysfunction, altered nuclear morphology, impaired nuclear membrane barrier and increased overall γH2AX signal; ^5^ here we add to the list the presence of chromatin remodeling and nucleolar stress. We further show, in non-myocytes, SAHFs, SASP, abundant p21 expression, and increased biological age. Just as these elements lead to (or accompany) neuronal degeneration, they may also contribute to the loss of cardiomyocytes in PKP2-deficient hearts. Though at first glance semantic, PKP2-ARVC shares an interesting feature in common with neurodegenerative diseases in that the primary insult to the molecular integrity of the cell is a path toward cell loss. This feature is shared with other cardiomyopathies (e.g., laminopathies) and it is different from other conditions (e.g., hypertrophic cardiomyopathies) in which the primary disruption causes a change in cellular physiology that, as an accumulated consequence, leads to a functional deficit in the heart.

### Limitations and future directions

It is important to note that our animal model does not reproduce all of the elements of PKP2-ARVC as a human disease. None of the experimental systems that have been utilized to study ACM reach that criterion. Yet though imperfect in many ways, the PKP2cKO model has served as a foundation of translational studies that have impacted PKP2-ARVC therapy in humans, such as the repurposing of flecainide as antiarrhythmic therapy^15,64,65^ and the preclinical studies for implementation of PKP2 gene therapy in human clinical trials. ^66,67^

The studies presented here provide a yet-untested path toward novel therapeutic strategies in ARVC. Indeed, cell senescence and neurodegenerative diseases are both conditions that have triggered extensive research as targets for therapy. Cell senescence associates with the arrest of cell division and as such, it is a desirable outcome in cancer therapy. ^68^ Yet, senolytic and senomorphic compounds have also been developed to tackle a variety of diseases such as idiopathic pulmonary fibrosis^69,70^, Alzheimer’s^71,72^ and accelerated aging syndromes^73^, to name a few, and remain to be tested for cardiac-related conditions. ^74^ Similarly, in view of the data presented here and in other studies, it may be reasonable to test, for conditions such as PKP2-ARVC, therapies aimed at ameliorating the progression of neurodegenerative diseases in a non-neurocentric way.

**In conclusion,** we have provided evidence indicating that loss of PKP2 expression only in myocytes is sufficient condition to induce a pro-inflammatory senescent phenotype in non-myocytes, and the premature aging of the heart. This represents, to our knowledge, the first experimental study that intersects the field of cellular aging and senescence, with that of desmosomal arrhythmogenic cardiomyopathies. Our results further highlight molecular parallelisms with neurodegenerative diseases, leading us to propose that PKP2-ARVC is a cardio-degenerative disease. Beyond semantics, this novel angle can permit pharmacological cross-references that can be of potential benefit to treat a disease that, at present, remains without a cure.

## CLINICAL PRESPECTIVES

Age is the leading risk factor for death. Previous studies have documented that molecular imbalances can accelerate the biological aging process, and have demonstrated that a senescent, pro-inflammatory phenotype has a negative impact on health. Here we expand medical knowledge by examining the molecular pathophysiology of arrhythmogenic right ventricular cardiomyopathy through the lens of premature aging and senescence, using an experimental model of Plakophilin-2 (PKP2) deficiency. Our studies show that PKP2 deficiency in cardiomyocytes is sufficient condition to provoke premature aging of cardiac cells and the pro-inflammatory, senescent phenotype of non-myocytes. A parallelism between the resulting molecular phenotype and that observed in neuro-degenerative diseases (e.g., Alzheimer’s, Parkinson’s, ALS) is discussed.

### Translational outlook

By their very nature, extrapolation of findings obtained through experimental models needs to be done with caution. Yet, within its limitations, the model utilized for this study has provided valuable preclinical information for therapeutic interventions. The findings presented here support the notion that pharmacological tools being developed to target senescent cells in other conditions such as Alzheimer’s disease could be applied to contain the release of harmful molecules from senescent cells in the cardiac niche, and that this approach can mitigate the progression of the disease. Future studies will be necessary to test the latter hypothesis.

## ACKNOWLEDGMENTS

We would like to acknowledge the following core facilities at the Division of Advanced Research Technologies, New York University Grossman School of Medicine, New York, NY, USA for their invaluable support: The Microscopy Laboratory [RRID:SCR_017934], the Experimental Pathology Research Laboratory [RRID:SCR_017928], and the Genome Technology Center [RRID:SCR_017929]. These facilities are partially supported by the Cancer Center Support Grant P30CA016087 at NYU Langone’s Laura and Isaac Perlmutter Cancer Center. Additionally, the Experimental Pathology Research Laboratory is supported by NIH S10 Shared Instrumentation Grant S10 OD021747.

## Abbreviations

ARVC: Arrhythmogenic Right Ventricular Cardiomyopathy
PKP2cKO: Cardiomyocyte-specific Conditional Knockout of Plakophilin-2
SAHF: Senescence-Associated Heterochromatin Foci
SASP: Senescence-Associated Secretory Phenotype
DDR: DNA Damage Response/Repair
Ex-SIM2: Expansion Micoroscopy-Structured Illumination Microscopy
TnI: Troponin I
LAD: Lamin-Associated Domain
LV: Left Ventricle
RV: Right Ventricle
NCS: Neocarzinostatin
TAM: Tamoxifen
Dpi: Days Post-Induction

## Notes

**Sources of funding** This work was partly supported by NIH grants R35HL160840 (MD) and RO1-HL169961A (EMS; MD), by American Heart Association 24SCEFIA1156995 (Dallas, TX, USA; IdeL) and by the Wilton W. Webster Fellowship in Electrophysiology from the Heart Rhythm Society (Washington, DC, USA; GB).

### Competing Interest Statement

The authors have declared no competing interest.

