## Supplemental appendix for "Cardiomyocyte-specific plakophilin-2 loss is sufficient to induce aging and senescence of nonmyocytes. Relevance to arrhythmogenic cardiomyopathy"

### SUPPLEMENTAL MATERIAL

### SUPPLEMENTAL FIGURES

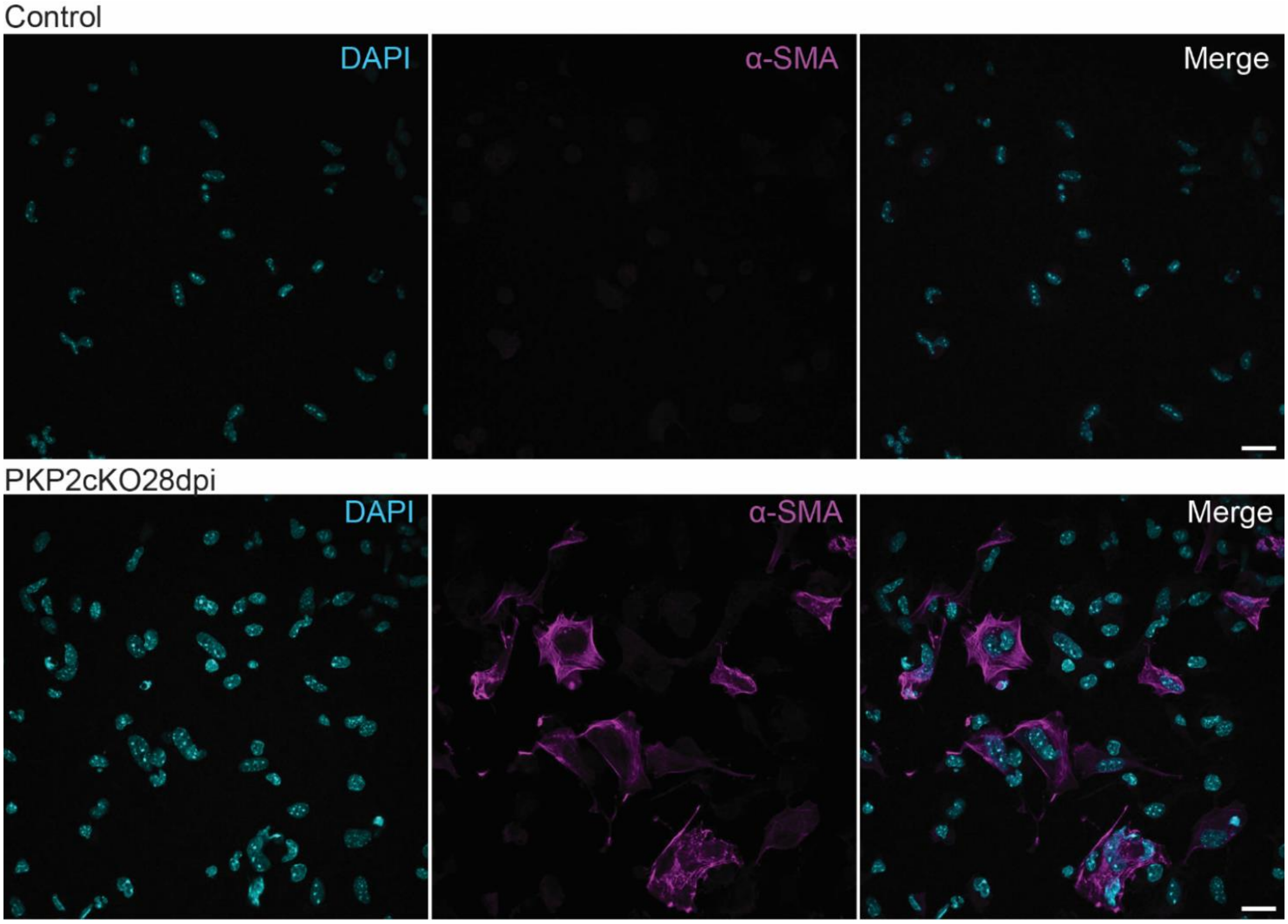


**Figure S1. Heterochromatin densities consistent with SAHFs in the nuclei of α-SMA-positive and of α-SMA-negative cells isolated from PKP2cKO hearts after separation from myocytes**. The image corresponds to that shown in Figure 1A, with the inclusion of α-SMA staining. Upper panels: Non-myocytes isolated from control hearts stained with DAPI (cyan) and α-SMA (magenta); merged image shown in the right panel. Lower panels: Same staining applied to non-myocytes from PKPcKO hearts at 28 dpi. Scale bar: 20 µm.

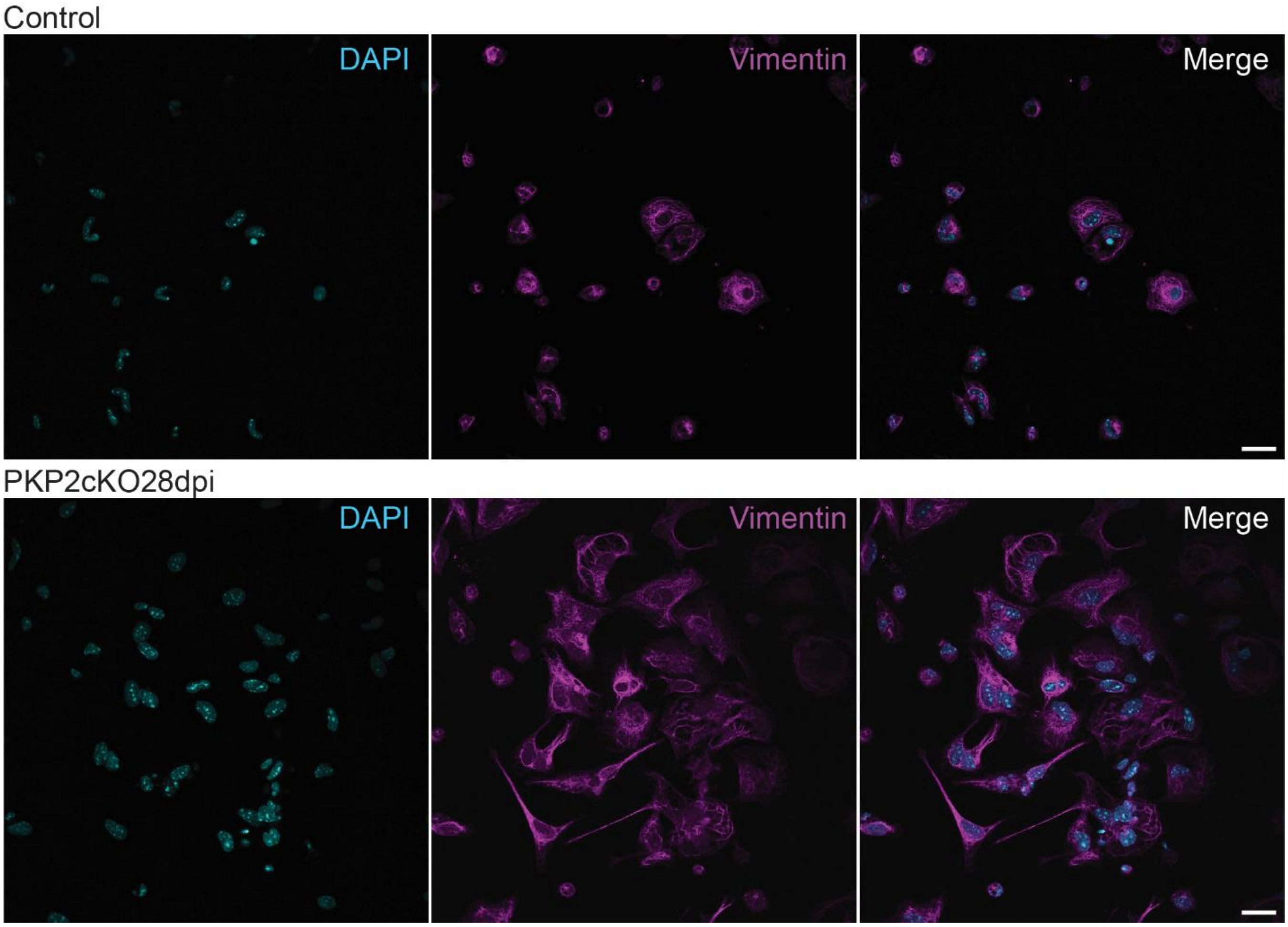


**Figure S2. Heterochromatin densities consistent with SAHFs in the nuclei of vimentin-positive cells isolated from PKP2cKO hearts after separation from myocytes**. Upper panels: Non-myocytes isolated from control hearts stained with DAPI (cyan) and Vimentin (magenta); merged image shown in the right panel. Lower panels: Same staining applied to non-myocytes from PKPcKO hearts at 28 dpi. Scale bar: 20 µm.

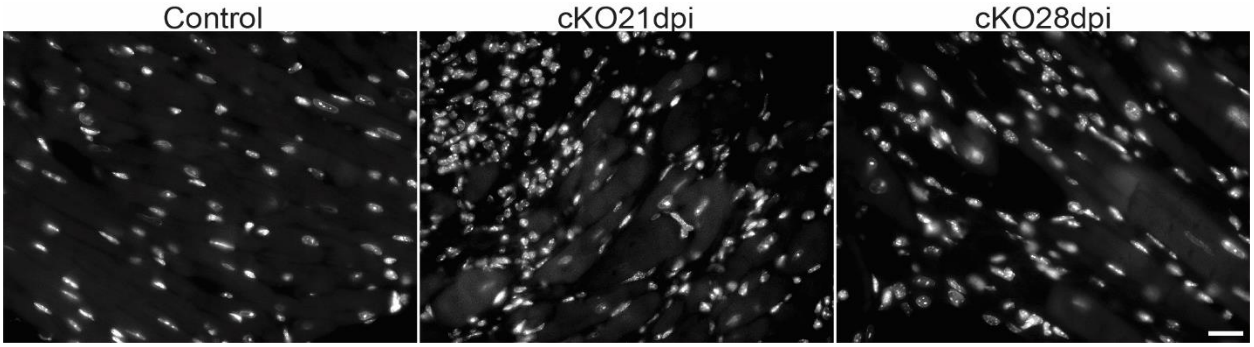


**Figure S3. Heterochromatin densities observed in PKP2cKO cardiac cells *in situ*.** Heart tissue sections from the were counterstained with DAPI. Left: Control; center: PKP2cKO 21 dpi; right: PKP2cKO 28 dpi. Scale bar: 20 µm.

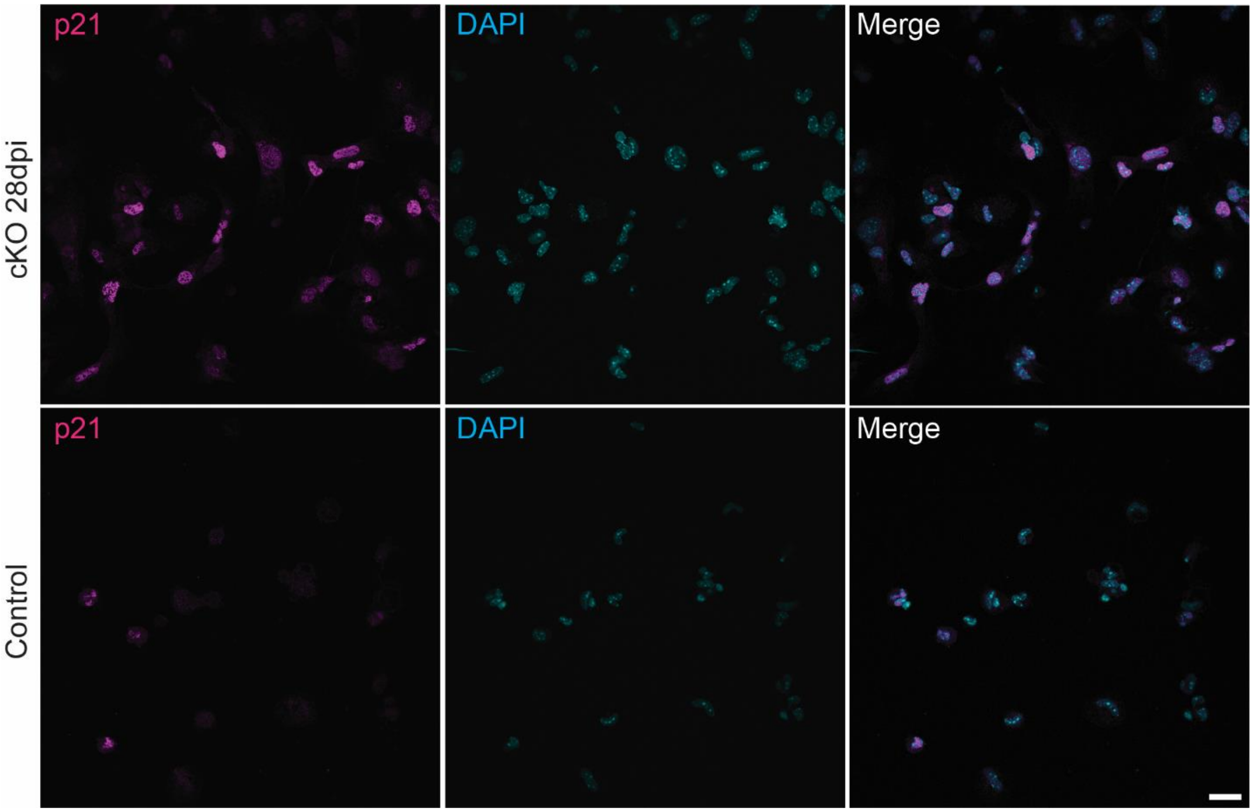


**Figure S4. p21 Expression in Non-Myocytes Isolated from PKP2cKO Hearts**. Representative images of non-myocyte nuclei isolated from control hearts (bottom) and PKP2cKO hearts 28 dpi (top). Left panels show p21 staining (magenta); center panels show DAPI staining (cyan); right panels display merged images. Scale bar: 20 µm.

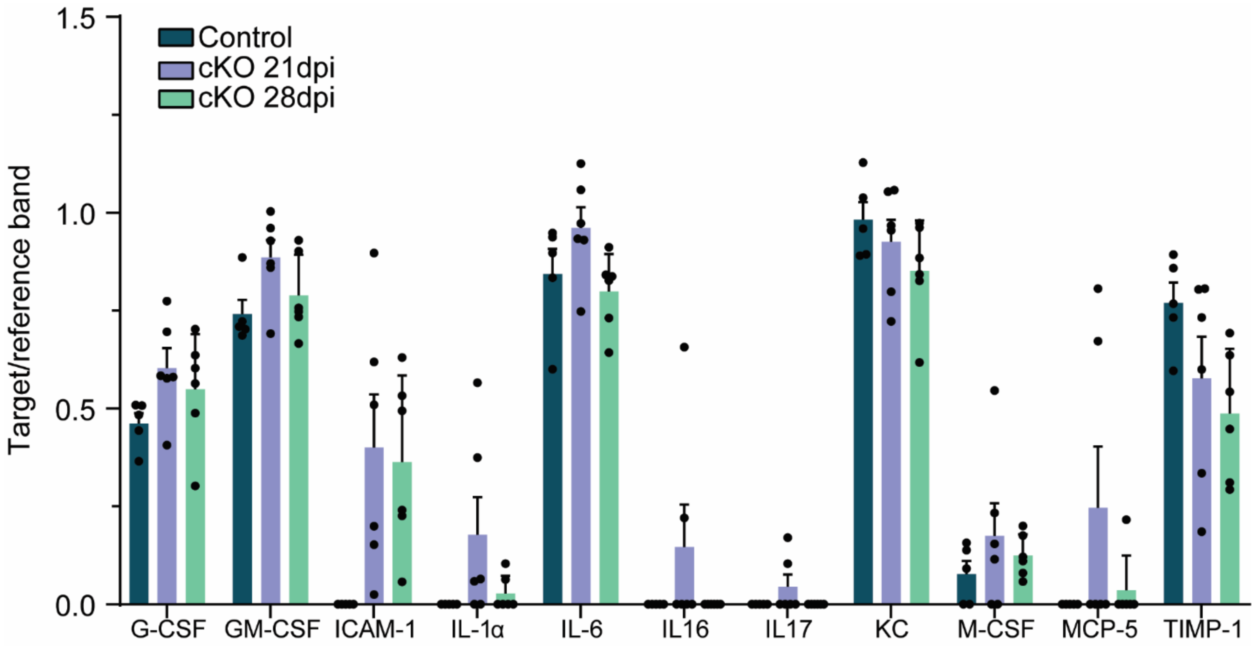


**Figure S5. Cytokine Profile of PKP2cKO Non-Myocyte Conditioned Media.** Bar graphs show density of the target signal relative to reference band, using a commercially available mouse cytokine array (see "Methods" for specifics). Data corresponds to targets detected in the conditioned media of PKP2cKO non-myocytes at least at one time point but not showing statistical significance from the control value. Data are shown as mean ± SD, each dot corresponds to a mouse. Normality was assessed using the Shapiro–Wilk test, and statistical testing was performed either by ANOVA or by a non-parametric test, as appropriate (see **Table S3**).

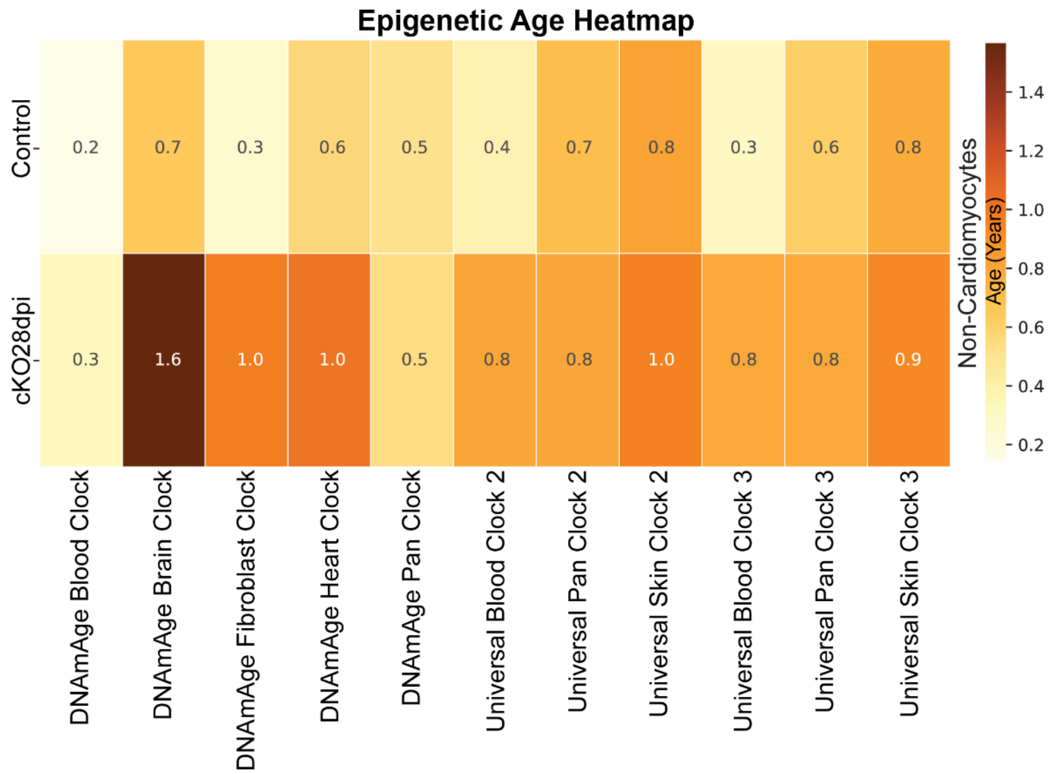


**Figure S6. Non-myocytes from PKP2cKO exhibit increased biological age across multiple clocks.** Heat map showing the predicted biological age (in years) of non-myocytes from control and PKP2cKO 28 dpi samples, based on CpG methylation profiles (epigenetic clocks). The numerical ages are reported within each square; darker color intensity indicates older biological age (see legend on the right). Though the estimated age of the control group varied, non-myocytes from PKP2cKO 28 dpi consistently showed increased biological age compared to controls across all clocks. Mouse-specific clocks (DNAmAge) [23] are shown first, followed by universal clocks that predict biological age across mammalian species and tissues. [24]

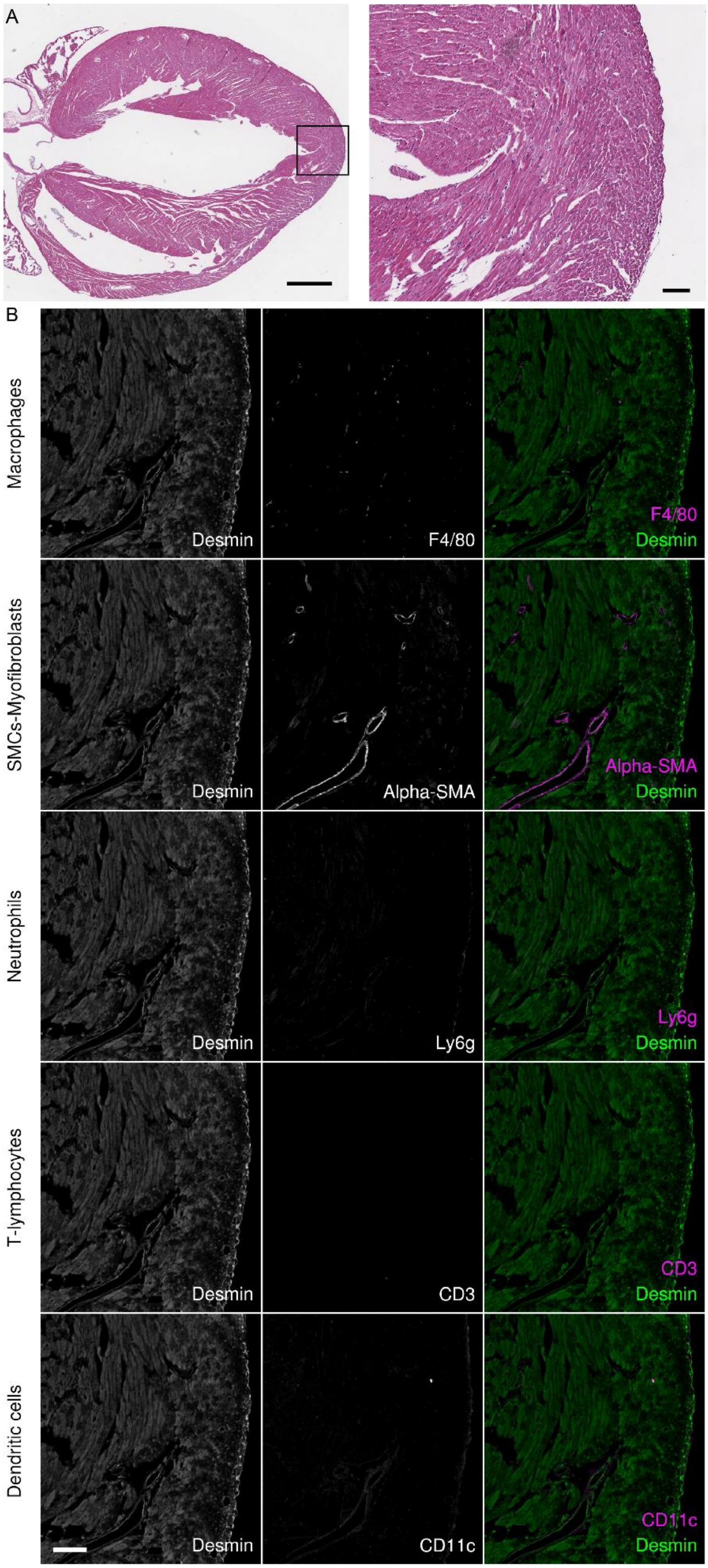

A
B
Macrophages
SMCs-Myofibroblasts
Neutrophils
T-lymphocytes
Dendritic cells
Desmin
F4/80
Alpha-SMA
Ly6g
CD3
CD11c

**Figure S7. Multiplex immunohistochemistry of a control heart.** (**A**) Hematoxylin and eosin (H&E) staining. Left: whole heart; right: magnified view of the area within the box. (**B**) Immunohistochemistry of the immediately subjacent section, corresponding to the region shown in A (right), using Opal 6-plex staining to label macrophages (F4/80), myofibroblasts (α-SMA), neutrophils (Ly6G), T lymphocytes (CD3), and dendritic cells (CD11c). Individual marker images are shown in the middle panels. Desmin staining (left panels) highlights myocytes as well as epicardial cells (desmin-positive, negative for all other markers). Merged images are shown in the right panels. Scale bars: 1 mm (A, left); 100 µm (A, right; B).

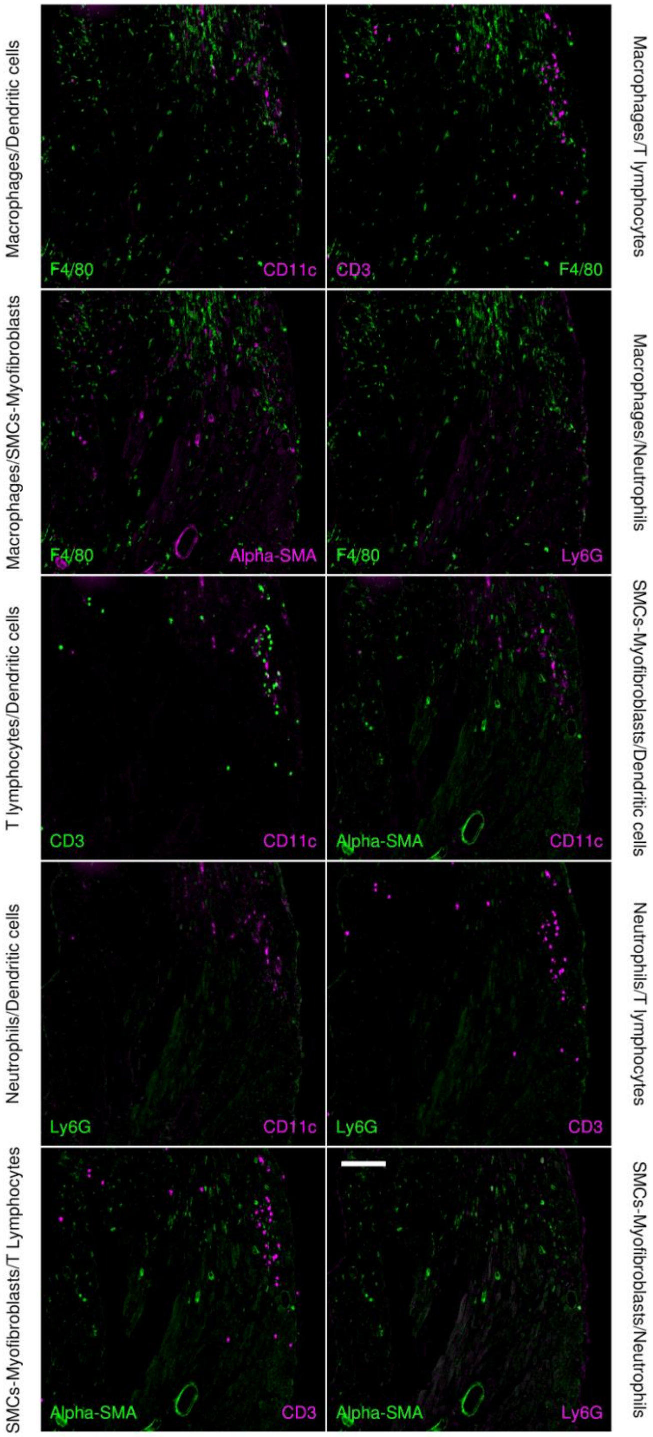


:

**Figure S8. Non-myocytes in PKP2cKO Hearts at 28 dpi.** Multiplex immunocytochemistry of non-myocyte populations in a PKP2cKO heart at 28dpi. Same sample as presented in Figure 2, here shows all pairwise combinations to emphasize the niches of cell populations and their intermingling. The specific antibodies/cell type studied are indicated within the figure. Scale bar: 100 µm.

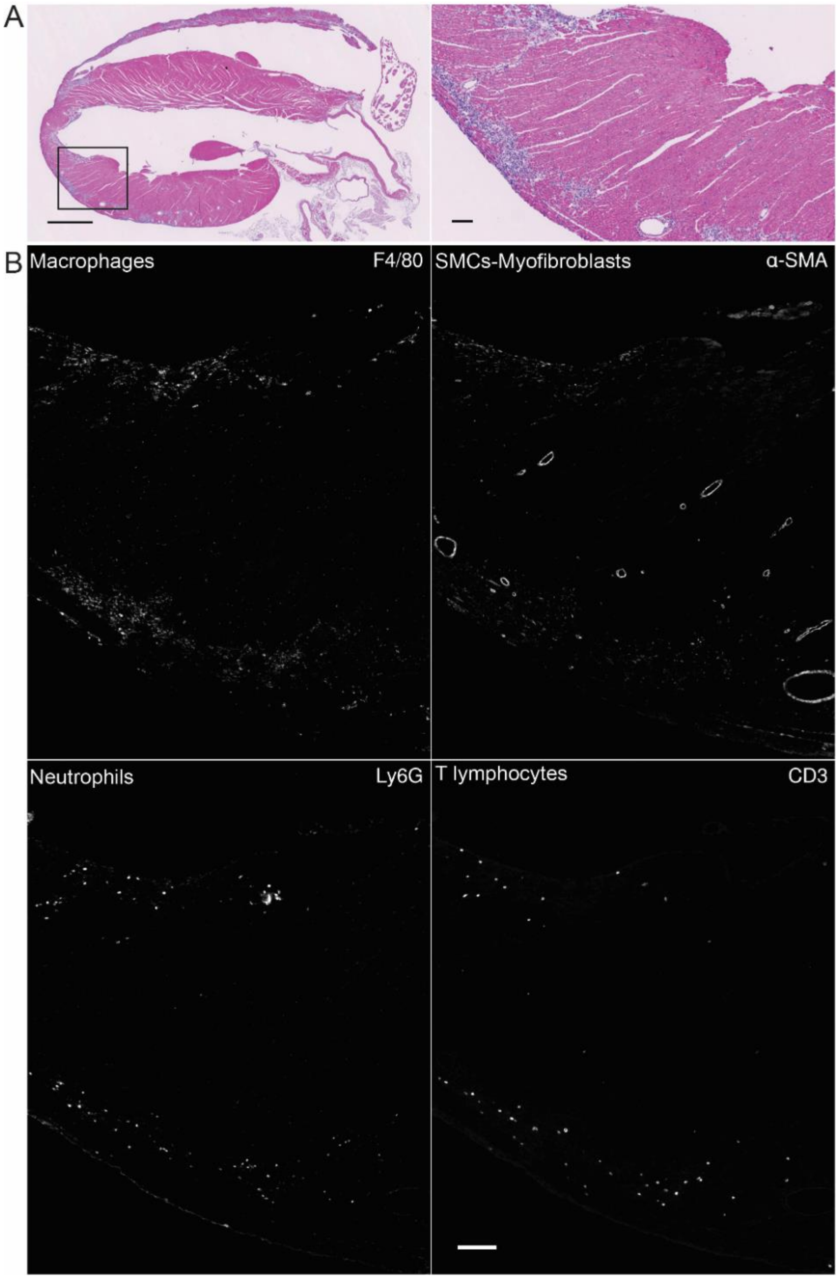


**Figure S9. Non-myocyte cells in PKP2cKO hearts at 21 dpi.** (**A**) Hematoxylin and eosin (H&E) staining of a PKP2cKO heart at 21 dpi in a four-chamber view. Panel on the right shows an enlarged view of the area indicated by the square in the left panel. (**B**) Immunostaining of the same region showed in the right of panel A, captured in a subjacent z-plane and labeled with four different antibodies. The identity of each non-myocyte cell type and the corresponding antibody are indicated in each image. Scale bars: 1 mm A left panel; 100 µm A right panel and panel B.

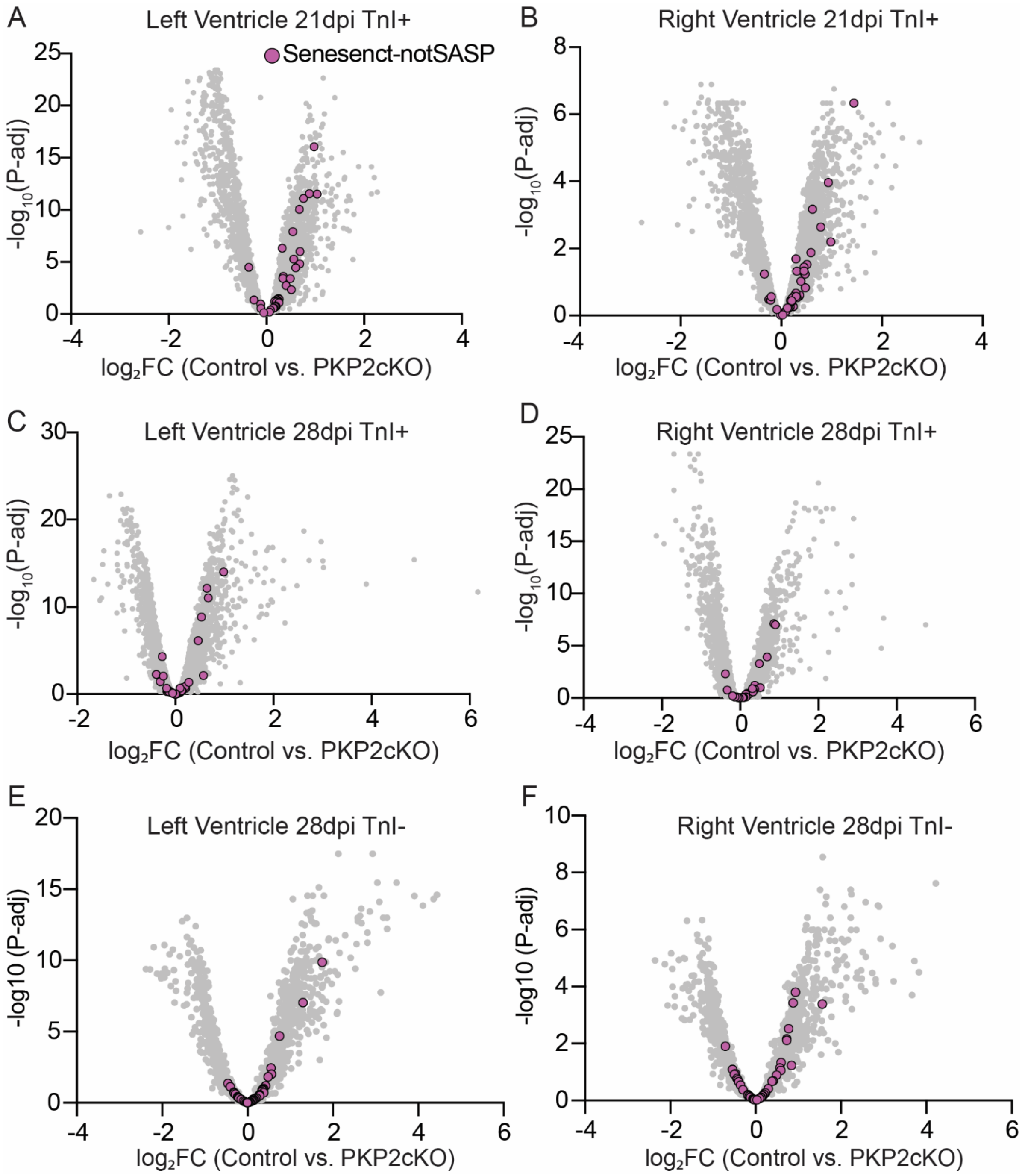


**Figure S10. Senescence-Related Transcripts (Excluding SASP) in PKP2cKO Hearts.** Volcano plots showing differential transcriptome analyses of cardiomyocyte-enriched areas ($TnI^{+}$) comparing Control vs. PKP2cKO hearts at 21 and 28 dpi: LV 21 dpi (**A**), RV 21 dpi (**B**), LV 28 dpi (**C**), and RV 28 dpi (**D**). (**E–F**) Volcano plots of $TnI^{-}$ areas comparing Control vs. PKP2cKO at 28 dpi in LV (**E**) and RV (**F**). Gray dots represent all differentially expressed transcripts, senescence-related transcripts (excluding SASP) are highlighted in purple. Note: for display purposes, the x- and y-axis ranges differ across the plots.

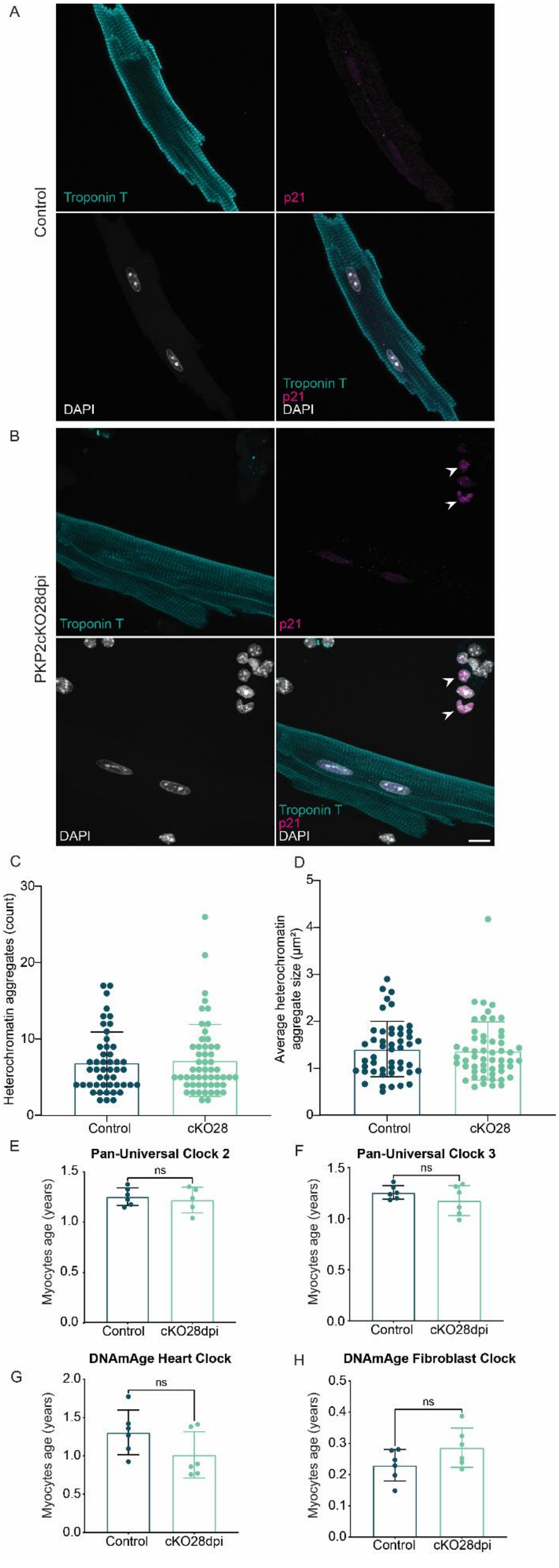
A
Control
Troponin T
p21
DAPI
Troponin T
p21
DAPI
B
PKP2cKO28dpi
Troponin T
p21
DAPI
Troponin T
p21
DAPI
C
Heterochromatin aggregates (count)
Control
cKO28
D
Average heterochromatin aggregate size (µm²)
Control
cKO28
E
Pan-Universal Clock 2
Myocytes age (years)
ns
Control
cKO28dpi
F
Pan-Universal Clock 3
Myocytes age (years)
ns
Control
cKO28dpi
G
DNAmAge Heart Clock
Myocytes age (years)
ns
Control
cKO28dpi
H
DNAmAge Fibroblast Clock
Myocytes age (years)
ns
Control
cKO28dpi

**Figure S11. Premature aging and senescence markers in cardiac cells. Panel A:** An isolated cardiomyocyte from a control heart stained for troponin T (cyan), p21 (magenta), and DAPI (gray). **Panel B**: Cells isolated from a PKP2cKO heart at 28 dpi. Same stainings as in A. For this experiment, myocytes and non-myocytes were kept in the same culture to allow comparison of phenotype between cell types. Notice the absence of p21 staining in the Troponin T-positive cell (i.e., a cardiomyocyte) and in contrast, the positive p21 staining in the non-myocyte cells indicated by the arrowheads in panel B. Scale bar: 10µm (applies to all panels) Heterochromatin densities consistent with SAHFs are also observed in the p21-labelled cells. **Panel C:** Quantification of heterochromatin aggregates in DAPI-labelled control and PKP2cKO cardiomyocytes at 28 dpi. **Panel D**: Average size of heterochromatin aggregates. Data are shown as mean ± SD, with each point representing one cell. For panels B-C, n/N (number of cells/Number of mice) are 48/3 and 54/3 for control and PKP2cKO 28 dpi, respectively. **Panels E-H**: predicted biological age (in years) of isolated cardiomyocytes from control and PKP2cKO 28 dpi samples, based on CpG methylation profiles (epigenetic clocks). Data are shown as mean ± SD, with each point representing one mouse (n=6 for each condition). In contrast to the results obtained from non-myocytes, there were no statistically significant differences between the groups.

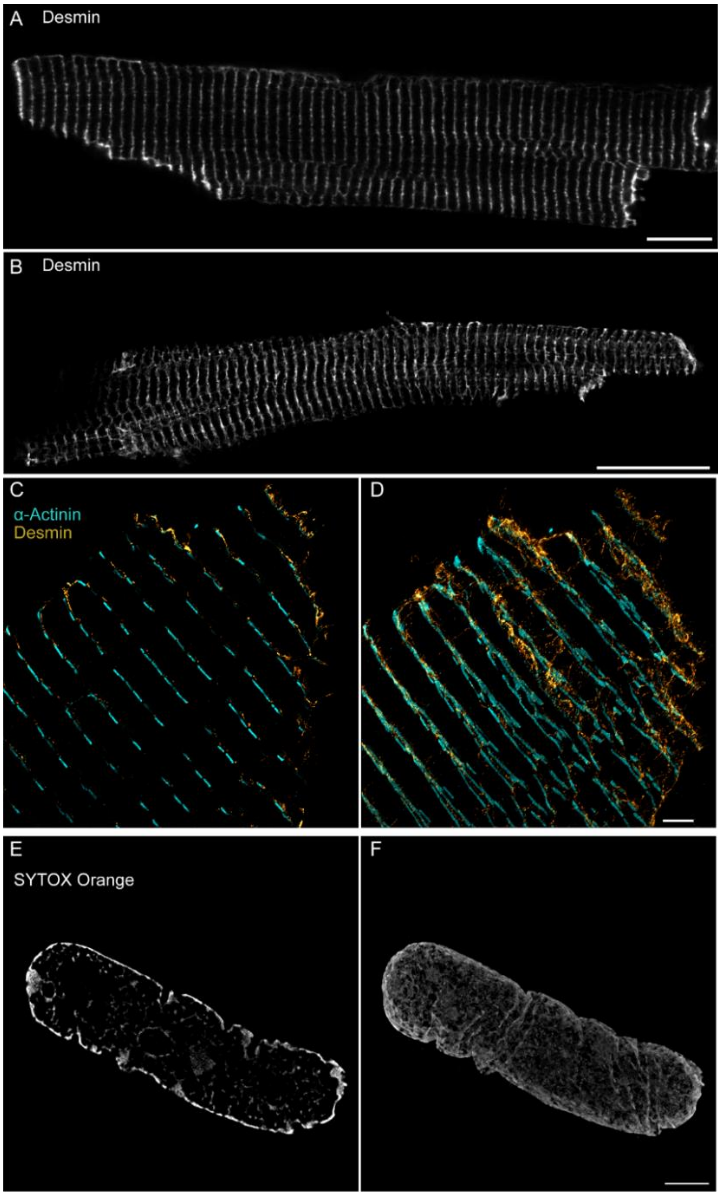


**Figure S12. Expansion and structure illumination microscopy applied to adult ventricular cardiomyocytes. A**) Representative image of a ventricular cardiomyocyte from a control mouse heart, stained for desmin. Image acquired using a Leica SP5 confocal microscope with a 63× oil immersion lens. Scale bar: 10 µm. **B**) Representative image of a separate cardiomyocyte processed with the expansion protocol, acquired using the same microscope with a 10× dry lens. Scale bar: 100 µm. Note the increase in overall cell size approaching 1 mm in length while maintaining clear sarcomeric organization. (**C–D**) Expanded cardiomyocyte stained with Desmin (gold) and α-actinin (cyan), imaged at the cell end using a Zeiss Elyra 7 microscope equipped for structured illumination microscopy and reconstructed with SIM². Panel C shows a single optical slice; panel D shows a Z-stack projection. Scale bar: 10 µm (applies to both panels). See also **Videos 1S and S2**. (**E–F**) Expanded and SIM² images of a nucleus from an adult murine ventricular myocyte stained with SYTOX Orange. Panel E shows a single Z-plane highlighting prominent peripheral heterochromatin and a central circular nucleolus. Panel F shows a 3D Z-stack reconstruction. Scale bar: 10 µm (applies to both panels). See also **Video S3**.

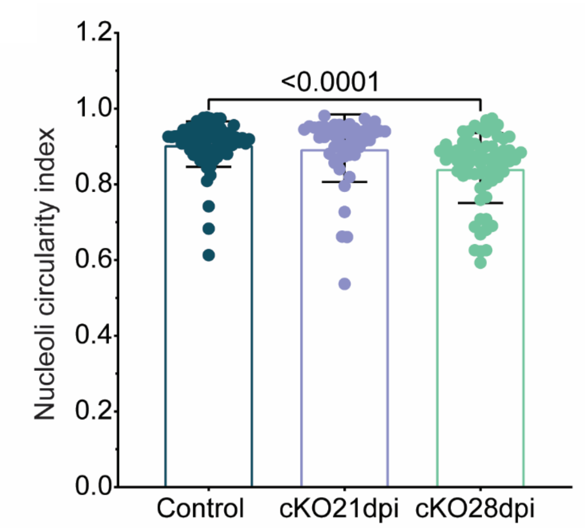


**Figure S13. Quantification of Nucleolar Morphology in PKP2cKO cardiac myocytes.** Bar graphs showing the nucleolar circularity index. Data are presented as mean ± SD, with each point representing one cell n/N=74/9, 46/5 and 64/7 for control, PKP2cKO 21 dpi and PKP2cKO 28 dpi respectively (n=number of cells; N=number of mice). Clustering was negligible (ICC < 5%, no superior multilevel fit).[20] Normality was assessed using the Shapiro–Wilk test and significance was detrmined by Kruskal–Wallis test followed by Dunn's multiple comparisons.

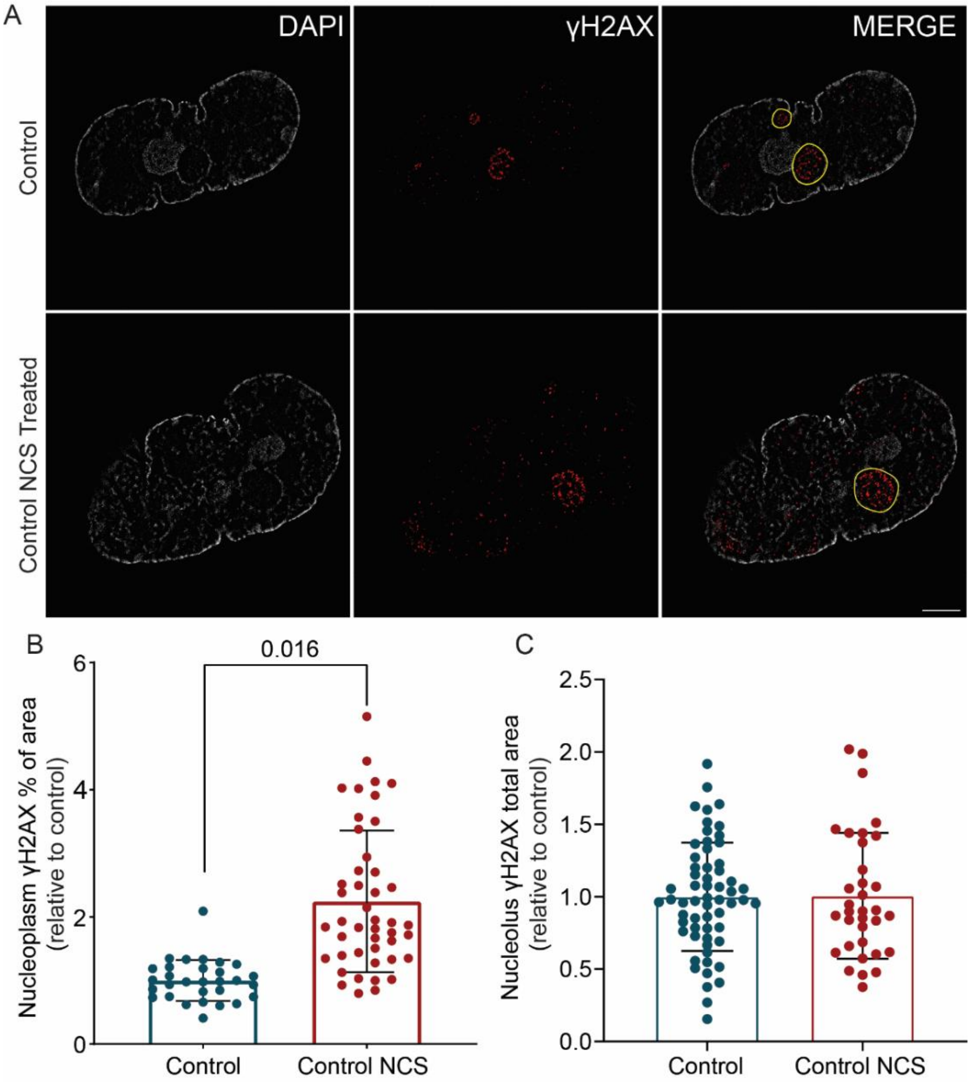


**Figure S14.Quantification of the γH2AX immunofluorescent signal elicited by NCS treatment in the nuclei of control cardiomyocytes.**

**A**) Representative images of a control nucleus (top) and a control nucleus treated with NCS (bottom), stained with DAPI (white) and γH2AX (red). Merged images are shown in the right panels; nucleoli are outlined in yellow to help identification. Scale bar: 300 nm. **B**) Bar graphs showing the percentage of γH2AX-positive pixels in the nucleoplasm (excluding nucleoli), normalized to the control average for each experiment. Data are presented as mean ± SD, with each point representing one cell. Control: 4 mice, 34 cells; Control + NCS: 4 mice, 46 cells. Clustering analysis revealed a high intraclass correlation (ICC = 33.5%) and a significantly better fit using a group-level model ($p < 0.0001$).[17] Statistical analysis was therefore performed accounting for clustering. **C**) Bar graphs showing the total γH2AX-positive area in the Nucleoli, normalized to the control average for each experiment. Data are shown as mean ± SD, with each point representing one cell. The control group is included for reference, as these data also appear in Figure 7. Control + NCS: 4 mice, 34 cells. Clustering analysis revealed a high intraclass correlation (ICC = 57.5%) and a significantly better fit using a group-level model ($p < 0.0001$).[17] Statistical analysis was therefore performed accounting for clustering.

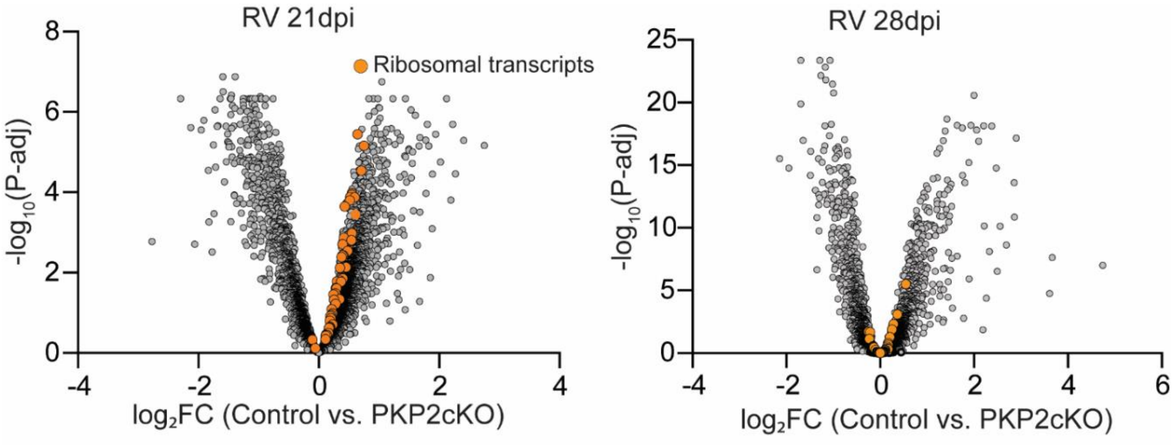


**Figure S15. Differential Expression of Ribosomal Protein Genes in samples collected from the right ventricle of PKP2cKO hearts.** Volcano plots showing differential transcriptome analysis of right ventricles from control vs. PKP2cKO mice at 21 (left) and 28 (right) dpi. Gray dots represent all detected transcripts; orange dots highlight transcripts from the Ribosomal Protein Gene Database.

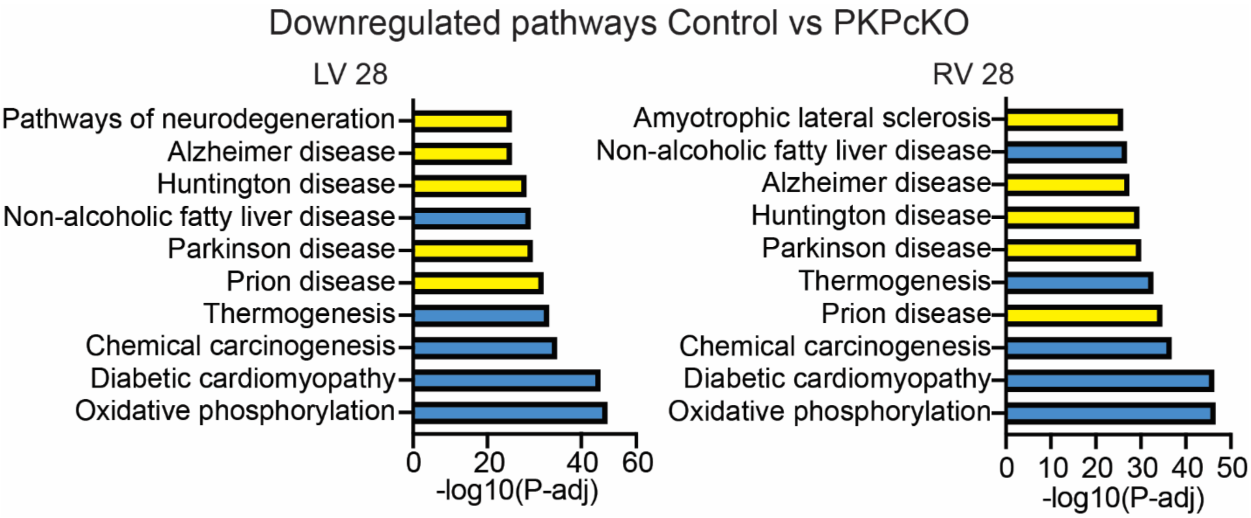


**Figure S16. Over-representation of Neurodegenerative Disease-Related Transcripts in PKP2cKO TNI- areas.** Kyoto Encyclopedia of Genes and Genomes (KEGG) pathway analysis of downregulated transcripts in control vs PKP2cKO LV and RV samples (left and right panels, respectively) collected at 28 dpi. The top 10 significant pathways are shown. Neurodegenerative disease pathways are highlighted in yellow. Thresholds of $|\log_2FC| > 0.5$ and adjusted $p < 0.005$ were applied.

### SUPPLEMENTAL TABLES

| Target | Host species | Supplier | Catalog number | Lot number | Concentration |
|---|---|---|---|---|---|
| γH2AX | Mouse | Sigma-Aldrich | Jbw301 | 3390229 | 1:25 |
| Fibrillarin | Rabbit | Abcam | AB5821 | 1050993-3 | 1:100 |
| UBF | Mouse | Santa Cruz Biotechnolgy | Sc-13125 | G2324 | 1:25 |
| Desmin | Rabbit | Scientific | PA516705 | ZG4414262 | 1:50 |
| α-actinin | Mouse | Sigma-Aldrich | A7811 | 314956 | 1:100 |

**Table S1**. Antibodies used for immunofluorescence of expanded cardiomyocytes, including target, host species, supplier, catalog number, lot number, and dilution.

| Antigen | Clone | Vendor, Cat.# | Retrieval | RRID | Blocker | Dilution Factor | Incubation Time [min] | Secondary or 2o HRP Polymer | Vendor/Cat # | Incubation Time [min] | Opal TSA Fluorophore | Vendor/Cat# | Fluorophore Dilution | Incubation Time [min] |
|---|---|---|---|---|---|---|---|---|---|---|---|---|---|---|
| **aSMA** | Poly | Abcam, ab5694 | ER1-20 | AB_2223021 | Leica Diluent | 200 | 30 | Rabbit-on -Rodent HRP polymer | Biocare/RMR622L | 10 | 520 | Akoya/FP1487001 KT | 150 | 10 mins |
| **Ly6g** | 1A8 (RUO) | BD, 551459 | ER1-60 | AB_394206 | Rodent Block M | 400 | 60 | Rat-1-step HRP polymer | Biocare/GHP516 | 10 | 570 | Akoya/FP1488001 KT | 150 | 10 mins |
| **CD3** | E4T1B | CST, 78588S | ER2-20 | AB_2889902 | Leica Diluent | 600 | 30 | Rabbit-on -Rodent HRP polymer | Biocare/RMR622L | 10 | 620 | Akoya/FP1495001 KT | 150 | 10 mins |
| **F480** | D2S9R | CST, 70076S | ER2-20 | AB_2799771 | Leica Diluent | 1500 | 30 | Rabbit-on -Rodent HRP polymer | Biocare/RMR622L | 10 | 480 | Akoya/FP1500001 KT | 200 | 10 mins |
| **CD11c** | D1V9Y | CST, 97585 | ER2-20 | AB_2800282 | Leica Diluent | 300 | 30 | Rabbit-on -Rodent HRP polymer | Biocare/RMR622L | 10 | 690 | Akoya/FP1497001 KT | 150 | 10 mins |
| **Desmin** | Poly | Invitrogen, PA5-115113 | ER2-20 | AB_2899749 | Leica Diluent | 1000 | 30 | Rabbit-on -Rodent HRP polymer | Biocare/RMR622L | 10 | 780 | Akoya/FP1501001[illegible] | 25 | 60 mins |

**Table S2.** Antibodies and specifications used for Opal 6-plex multiplex immunohistochemistry of heart tissue.

| Cytokine/Chemokine | mean value CTRL | mean value cKO21 | mean value cKO28 | p-value CTRL vs cKO21 | p-value CTRL vs cKO28 | pass Normality | statistical test |
|---|---|---|---|---|---|---|---|
| G-CSF | 0.4616 | 0.6029 | 0.5492 | ns | ns | Yes | Ordinary one-way ANOVA/ Dunnett's multiple comparisons test |
| GM-CSF | 0.7222 | 0.8702 | 0.7891 | ns | ns | No | Kruskal-Wallis test/Dunn's multiple comparisons test |
| ICAM1 | N/A | 0.4003 | 0.3636 | 0.0116 | 0.0082 | No | Kruskal-Wallis test/Dunn's multiple comparisons test |
| IL -1α | N/A | 0.1773 | 0.0278 | ns | ns | No | Kruskal-Wallis test/Dunn's multiple comparisons test |
| IL-1ra | 0.3625 | 0.6957 | 0.6811 | 0.0099 | 0.0131 | Yes | Ordinary one-way ANOVA/ Dunnett's multiple comparisons test |
| IL-6 | 0.8337 | 0.9613 | 0.7985 | ns | ns | Yes | Ordinary one-way ANOVA/ Dunnett's multiple comparisons test |
| IL-10 | N/A | 0.2432 | 0.1477 | ns | 0.046 | No | Kruskal-Wallis test/Dunn's multiple comparisons test |
| IL-16 | N/A | 0.1463 | N/A | ns | ns | No | Kruskal-Wallis test/Dunn's multiple comparisons test |
| IL-17 | N/A | 0.0457 | N/A | ns | ns | No | Kruskal-Wallis test/Dunn's multiple comparisons test |
| IP-10 | 0.1147 | 0.7089 | 0.693 | 0.0108 | 0.0108 | Yes | Ordinary one-way ANOVA/ Dunnett's multiple comparisons test |
| KC | 0.9819 | 0.9256 | 0.8259 | ns | ns | Yes | Ordinary one-way ANOVA/ Dunnett's multiple comparisons test |
| M-CSF | 0.0773 | 0.1746 | 0.1102 | ns | ns | Yes | Ordinary one-way ANOVA/ Dunnett's multiple comparisons test |
| JE | 0.6069 | 0.3254 | 0.3276 | 0.0176 | 0.016 | Yes | Ordinary one-way ANOVA/ Dunnett's multiple comparisons test |
| MCP-5 | N/A | 0.2463 | 0.036 | ns | ns | No | Kruskal-Wallis test/Dunn's multiple comparisons test |
| MIP-1 α | 0.6656 | 0.9913 | 0.8992 | 0.0019 | 0.0185 | Yes | Ordinary one-way ANOVA/ Dunnett's multiple comparisons test |
| MIP-1β | 0.0969 | 0.7113 | 0.6795 | <0.0001 | <0.0001 | Yes | Ordinary one-way ANOVA/ Dunnett's multiple comparisons test |
| MIP-2 | 0.715 | 0.4148 | 0.4012 | 0.0286 | 0.0222 | Yes | Ordinary one-way ANOVA/ Dunnett's multiple comparisons test |
| RANTES | 0.0689 | 0.8128 | 0.7922 | 0.0091 | 0.0128 | Yes | Ordinary one-way ANOVA/ Dunnett's multiple comparisons test |
| SDF-1 | 0.0641 | 0.214 | 0.1673 | 0.0488 | ns | Yes | Ordinary one-way ANOVA/ Dunnett's multiple comparisons test |
| TIMP-1 | 0.7673 | 0.577 | 0.4869 | ns | ns | Yes | Ordinary one-way ANOVA/ Dunnett's multiple comparisons test |
| TNF-α | 0.1312 | 0.5401 | 0.3355 | 0.0087 | ns | Yes | Ordinary one-way ANOVA/ Dunnett's multiple comparisons test |
| BLC | - | - | - | - | - | - | - |
| C5/C5a | - | - | - | - | - | - | - |
| I-309 | - | - | - | - | - | - | - |
| Eotaxin | - | - | - | - | - | - | - |
| IFN-γ | - | - | - | - | - | - | - |
| IL-1β | - | - | - | - | - | - | - |
| IL-2 | - | - | - | - | - | - | - |
| IL-3 | - | - | - | - | - | - | - |
| IL-4 | - | - | - | - | - | - | - |
| IL-5 | - | - | - | - | - | - | - |
| IL-7 | - | - | - | - | - | - | - |
| IL-13 | - | - | - | - | - | - | - |
| IL-12p70 | - | - | - | - | - | - | - |
| IL-23 | - | - | - | - | - | - | - |
| IL-27 | - | - | - | - | - | - | - |
| I-TAC | - | - | - | - | - | - | - |
| MIG | - | - | - | - | - | - | - |
| TARC | - | - | - | - | - | - | - |
| TREM-1 | - | - | - | - | - | - | - |

**Table S3.** List of chemokines and cytokines included in the Proteome Profiler Mouse Cytokine Array Kit, Panel A (R&D Systems, Cat# ARY006). Mean densitometry values, normalized to the reference dots, are reported for detected molecules, entries without values indicate cytokines not detected in the samples. Normality was assessed using the Shapiro–Wilk test. Statistical significance was evaluated by multiple comparison testing, with p-values reported for control versus PKP2cKO21 and control versus PKP2cKO28 dpi.

**Table S4.** Differential transcriptomic analysis of $TnI^+$ areas across four conditions: control versus PKP2cKO21 (LV and RV) and control versus PKP2cKO28 (LV and RV). For each transcript and comparison, the adjusted p-value and log fold change are reported. Senescence-associated secretory phenotype (SASP) associated transcripts and other senescence-associated transcripts, as defined by Saul et al.,[22] are indicated within the main list.

**Table S5.** Differential transcriptomic analysis of $TnI^-$ areas comparing control versus PKP2cKO28 dpi. Results are provided separately for the LV and RV. For each transcript, the adjusted p-value and log fold change are reported. SASP and other senescence-associated transcripts, as defined by Saul et al.,[22] are indicated within the lists.

**Table S6.** List of genes represented in the Cartesian plot shown in Figure 3G. Genes present in each quadrant are reported (named Quadrant 1, 2, 3, and 4). For each transcript, the adjusted p-value and log fold change from the differential analysis of PKP2cKO28 versus control in LV and RV $TnI^+$ and $TnI^-$ regions are reported.

**Table S7.** Ribosomal transcripts identified in $TnI^+$ regions based on the Ribosomal Protein Gene Database.[34] For each transcript, the adjusted p-value and log fold change are reported for control versus PKP2cKO21 (LV, RV) and control versus PKP2cKO28 (LV, RV).

**Table S8.** Proteins dysregulated in control versus PKP2cKO hearts at 21 dpi (Pérez-Hernández et al.)[5] that overlap with proteins dysregulated during aging (6 vs 30 months; Takasugi et al.)[35]. Proteins with an adjusted p-value < 0.05 are reported. For each protein, the log fold change and adjusted p-value are presented.

**Table S9.** Transcripts corresponding to markers of neurodegenerative disorders (Alzheimer's disease, Parkinson's disease, and ALS; DISGENET).[37] Sheet 1 lists all identified transcripts, with log fold change and adjusted p-value reported for $TnI^+$ regions across four conditions: control versus PKP2cKO21 (LV and RV) and control versus PKP2cKO28 (LV and RV). Additional sheets report transcripts significantly dysregulated (adjusted p-value < 0.05) in only one condition (LV or RV at 21 dpi or 28 dpi) and transcripts shared across specific combinations of conditions, up to those found in all four datasets.

#### SUPPLEMENTAL VIDEOS

**Video S1.** Expanded cardiomyocyte stained with Desmin (gold) and α-actinin (cyan), imaged at the cell end using a Zeiss Elyra 7 microscope with structured illumination microscopy and reconstructed with SIM². The video displays the Z-stack plane by plane, capturing the central portion of the myocyte, and corresponds to panels C–D of Figure S8.

**Video S2.** Single Z-line from and expanded cardiomyocyte stained with Desmin (green) and α-actinin (red), imaged using a Zeiss Elyra 7 microscope with structured illumination microscopy and reconstructed with SIM². The video shows a 360° rotation along the x-axis of a Z-line from the same myocyte showed in to panels C–D of Figure S8.

**Video S3**. Expanded cardiomyocyte nucleus stained with SYTOX Orange, imaged using a Zeiss Elyra 7 microscope with structured illumination microscopy and reconstructed with SIM². The video displays the Z-stack plane by plane of a nucleus from an adult murine ventricular myocyte, corresponding to panels E–F of Figure S8.

**Video S4.** Nucleolus from a control cardiomyocyte stained for γH2AX (gold), fibrillarin (cyan), and chromatin marked by DAPI (white), imaged using a Zeiss Elyra 7 microscope with structured illumination microscopy and reconstructed with SIM². The video displays the Z-stack plane by plane, corresponding to the upper panel of Figure 5A.

**Video S5.** Nucleolus from a PKP2cKO cardiomyocyte (28 dpi) stained for γH2AX (gold), fibrillarin (cyan), and chromatin marked by DAPI (white), imaged using a Zeiss Elyra 7 microscope with structured illumination microscopy and reconstructed with SIM². The video displays the Z-stack plane by plane, corresponding to the lower panel of Figure 5A.

### EXTENDED METHODS

#### Animal model

Animal experiments were performed in 3–6-month-old male and female mice expressing a cardiomyocyte-specific, tamoxifen (TAM)-activated, deletion of *Pkp2* (PKP2cKO). Cre-negative, flox-positive, TAM-injected age and sex matched mice were used as controls. [15] PKP2cKO mice were used at 21 days post injection (21 dpi) of TAM, a time point at which an arrhythmogenic cardiomyopathy of right ventricular predominance can be observed and at 28 dpi, when left ventricular function is also compromised. [15] Procedures conformed with the Guide for Care and Use of Laboratory Animals of the National Institutes of Health and were approved by the New York University Institutional Animal Care and Use Committee (IA16-01021).

**Isolation of Murine Ventricular Myocytes and Non-Myocytes**

Murine ventricular myocytes and non-myocytes were obtained by enzymatic dissociation following standard procedures. [18] Briefly, mice were injected with 0.2 ml heparin (500 IU ml−1 intraperitoneally; Sigma Aldrich, Cat# H3393) 20 min before heart excision and anesthetized by inhalation of 100% CO2. Deep anesthesia was confirmed by lack of response to otherwise painful stimuli and cervical dislocation was performed after anesthesia to ensure the death of the animal. Hearts were quickly removed from the chest and placed in a Langendorff column. For cell dissociation, isolated hearts were perfused at 3 ml/min at 37°C with perfusion buffer containing (in mmol/l): 113 NaCl, 4.7 KCl, 1.2 $MgSO_4$, 0.6 $Na_2HPO_4$, 0.6 $KH_2PO_4$, 12 $NaHCO_3$, 10 $KHCO_3$, 10 HEPES, 5.5 glucose, and 30 taurine (pH 7.40 with NaOH). The hearts were then perfused with the same buffer supplemented with collagenase type II (600 U/ml; Worthington, Cat# LS004177) and 12.5 µM $CaCl_2$ for 15 minutes. The ventricles were minced; cardiac cells were suspended in stop buffer (perfusion buffer with 12.5 µM CaCl2 and 5% Fetal Bovine Serum [FBS; Gibco]) and passed through a 100-µm filter. Myocytes and non-myocytes were separated by differential sedimentation, using a modified protocol from Ackers-Johnson et al. [75] The cell suspension in stop buffer was first centrifuged at 30 × g for 3 minutes to pellet the myocytes. The myocyte pellet was then washed in perfusion buffer and centrifuged a second time to remove residual non-myocytes. The combined supernatants, enriched in non-myocytes, were collected and centrifuged at 300 × g for 5 minutes to obtain the non-myocytes fraction.

**Non-Myocyte Culture and Cytokine array**

Each non-myocyte cell pellet was resuspended in 1.8 mL of DMEM (Gibco, Cat# 10-013-CV) supplemented with 10% FBS and 1% Penicillin-Streptomycin (Gibco). The suspension was plated into six wells of a 24-well plate (300 µL per well), with glass coverslips placed in half of the wells for subsequent fixation and immunostaining. Cells were incubated at 37 °C in 5% $CO_2$ for 4 hours, then washed twice with PBS 1× to remove debris. Cells on coverslips were immediately fixed in 4% paraformaldehyde (PFA; Electron Microscopy Sciences) in PBS 1X for 10 minutes and stored in PBS 1x. Fresh DMEM (600 µL) was added to wells without coverslips. After 24 hours, the conditioned medium was collected, centrifuged at 200 × g for 10 minutes at 4 °C to remove cellular debris, and the clarified supernatants were pooled for subsequent cytokine analysis. Membranes from the Proteome Profiler Mouse Cytokine Array Kit, Panel A (R&D Systems, Cat# ARY006), were incubated with 1 mL of the pooled supernatant, and the assay was performed according to the manufacturer's instructions.

**Confocal Immunofluorescence of Myocytes and Non-Myocytes**

Freshly isolated ventricular myocytes were plated on 13 mm glass coverslips coated with laminin (1:10 dilution; Corning) and fixed for 15 minutes in 4% PFA in PBS 1X. Non-myocytes were fixed as described above. For detection of p21 expression, and for detection of senescence-associated heterochromatin foci, myocytes and non-myocytes fixed on glass coverslips were permeabilized for 10 minutes with 0.5% Triton X-100 (Sigma Adrich) and then blocked for 1 hour at room temperature in blocking buffer, which consisted of PBS 1X containing 0.02% Tween 20 (Sigma Aldrich) and 5% bovine serum albumin (BSA) Fraction V (Roche). Cells were incubated overnight at 4 °C with anti-p21 (1:200; Abcam, Cat# ab188224). Myocytes were co-incubated with anti–troponin T (1:200; Abcam, Cat# ab8295). After three washes in wash buffer, cells were incubated for 1 hour at room temperature with the following secondary antibodies: Alexa Fluor 488 goat anti-rabbit (1:300; Invitrogen, Cat# A11008) for non-myocytes, and for myocytes Alexa Fluor 647 goat anti-rabbit (1:500; Invitrogen, Cat# A21244) together with Alexa Fluor 568 goat anti-mouse (1:500; Invitrogen, Cat# A11031). Following three washes with PBS 1×, nuclei were counterstained with DAPI (1 μg/mL; Invitrogen, Cat# D1306). Coverslips were mounted onto glass slides using ProLong™ Gold Antifade Mountant (Invitrogen, Cat# P36930) and allowed to dry overnight at room temperature. Imaging was performed using a Leica TCS SP5 confocal microscope equipped with a 63×/1.2 oil-immersion objective. Images of non-myocytes were acquired as single z-plane sections while myocytes were acquired as z-stacks.

Image analysis was performed using ImageJ. Nuclei were delineated as individual Regions of Interest (ROIs) and used to measure nuclear areas of non-myocytes. An automatic fluorescence intensity threshold was applied, and only brighter DAPI-stained pixels, distinguishable from uniformly stained nuclei, were classified as heterochromatic foci (defined as SAHF in non-myocytes). SAHF above the defined threshold were manually counted by two independent investigators.

**DNA Methylation Profile and Epigenetic Clock analysis of myocytes and non-myocytes**

Myocyte and non-myocyte pellets obtained as described above were flash-frozen in liquid nitrogen. Genomic DNA was subsequently extracted using the DNeasy Blood & Tissue Kit (Qiagen, cat. 69504) and shipped to the Clock Foundation (Los Angeles, CA) for DNA methylation profiling and biological age predictions through epigenetic clocks. Methylation was measured across the genome in 12 mouse samples from myocytes and 12 samples from non-myocytes across two conditions (Control and PKP2cKO 28 dpi; n = 6 per group). Upon receipt, the Clock Foundation performed quality control, including quantification and integrity assessment, followed by bisulfite conversion using the Zymo EZ DNA Methylation Kit (Zymo Research, Cat# D5001), according to the manufacturer's protocol. DNA methylation was measured using the custom Illumina 320K mouse array, which interrogates over 285,000 CpG sites across the mouse genome. [19] Arrays were scanned using the Illumina iScan System, and raw IDAT files were generated. Data were processed and normalized using the SeSAMe R package, [76] resulting in beta values and associated detection p-values for each probe. Beta values represent the ratio of methylated probe fluorescence intensity to the total signal (sum of methylated and unmethylated probe intensities plus a constant), ranging from 0 (fully unmethylated) to 1 (fully methylated). [77] For downstream predictive analysis using reduced feature sets, the 320K array data were downsampled to a 40K subset. This was achieved by selecting probes that align between both arrays with a Pearson correlation coefficient exceeding 0.8. For any probes missing in the reduced dataset, gold median values were imputed based on a large mouse reference dataset curated by the Clock Foundation. Measures of epigenetic aging were derived using universal pan-tissue clock, applicable across mammalian species, and mouse-specific pan-tissue and tissue-specific clocks. [23,24]

**Chemical induction of DDR**

As a point of reference regarding the characteristics of the DDR signal in PKP2cKO myocytes, separate experiments were conducted in cardiomyocytes isolated from control mice and treated with 60 ng/mL of Neocarzinostatin (NCS; Sigma-Aldrich, Cat# N9162) for 30 minutes at 37°C.

**Expansion microscopy and Structured Illumination (Ex-SIM²)**

Following fixation, myocytes were permeabilized for 15 minutes with 0.5% Triton X-100 (Sigma Adrich) and then blocked for 1 hour at room temperature in blocking buffer (described above). Primary antibodies (listed in **Table S1**) in blocking buffer were incubated overnight at 4°C. After three washes with wash buffer (0.02% Tween 20 in PBS 1X), cells were incubated for 1 hour at room temperature with secondary antibodies diluted 1:100 in blocking buffer: Alexa Fluor 488 goat anti-rabbit (Invitrogen, Cat# A11008, Lot# 2690601) and Biotium 568 anti-mouse (Cat# 20100, Lot# 22C0421). Following three washes with wash buffer, the cross-linking reagent Acryloyl-X SE (ACX; Invitrogen; 1 mg/mL in PBS 1X) was applied overnight at room temperature to ensure protein anchoring to the hydrogel. After ACX treatment, 2 washes of 15 min with PBS 1X were performed followed by gelation as previously described. [26] Briefly, 13 mm silicone isolation chambers were filled with 100 µL of gelation mix[26], and the coverslips were placed on top of the gel drop with the cell side in contact with the gel. The assembled gelation chambers were incubated for 1 hour at 37°C. Following gelation, the coverslips with the polymerized gels on top were digested for 4 hours at 37°C with proteinase K (8 U/mL; ThermoFisher) in digestion buffer. [26] During digestion, nuclei were counterstained with either DAPI 1mg/mL (Invitrogen, Cat# D1306) 1:200 or SYTOX™ Orange (ThermoFisher, Cat# S11368) 0.25µM in digestion buffer. Following digestion, the gels spontaneously detached from the coverslips and were expanded in ultra-pure water. The water was changed at least once after

30 minutes, and the gels were left overnight at room temperature to allow full expansion. The expanded gels were carefully cut using glass slides and transferred to 35 mm glass-bottom dishes (Mattek, Cat# P35G-1.5-20-C) coated with poly-L-lysine overnight (Sigma Aldrich, Cat# P8920). The expansion factor was 6.4X. All distances of a value "Y" obtained from expanded cells are reported as corrected for expansion factor (i.e., Y/6.4). Unless specifically noted, all imaging of expanded samples in this study was performed using a Zeiss Elyra 7 structured illumination microscope equipped with a C-Apochromatic 63x/1.4 water objective and lattice $SIM^2$ technology. The images were acquired as a z-stack with a step size of 0.03 μm. The system allowed image reconstruction of large areas in the XY plane, as well as along the z axis, resulting in a three-dimensional view of the structures.

Optical resolution of images collected through structured illumination is estimated at approximately 60 nm. [57] Considering that distances expanded by a factor of ~6.4X, we calculated our optical resolution to be well below the 40 nm range. For image analysis, ROIs were manually drawn in ImageJ following the contours of the heterochromatic ring surrounding the nucleoli and used to measure nucleoli area and circularity index. The total area of positive pixels within each ROI was determined using an automatic fluorescence intensity threshold to distinguish positive from negative pixels. For the measurement of heterochromatin at the nuclear lamina (LAD), two ROIs were defined: one tracing the dense heterochromatic line at the nuclear periphery and another encompassing the entire nucleus. The area of heterochromatin above the fluorescence intensity threshold within the peripheral ROI was expressed relative to the total heterochromatin area of the nucleus. Results were then normalized to the average values of the corresponding control samples processed in parallel to PKP2cKO.

**Spatial Transcriptomics via NanoString GeoMx Digital Spatial Profiling**

The transcriptomics study was conducted on heart tissue from 6 PKP2cKO mice 21 days post-TAM, 6 PKP2cKO mice 28 days postTAM, and 12 Cre-negative controls matched to the PKP2cKO mice. Formalin-fixed and paraffin embedded tissue was sectioned along the long axis to render a four-chamber view. Sections were immunostained for Troponin I (TnI), as a myocyte marker. Areas in the free wall of either the RV or the LV that were populated densely by myocytes (as visualized by fluorescence microscopy) were marked as regions of interest (ROIs) using the Nanostring GeoMx Digital Spatial Profiler. [78] Five µm sections were prepared, thoroughly cleaning the microtome and recovering sections in RNAse free water to avoid introducing RNAses and decrease the time between sectioning and slide preparation. After sectioning, slides were stored momentarily at 4° C in a sealed box with silica pouches to prevent humidity and condensation.

Semi-automated slide preparation was performed according to the NanoString® standard of practice for NGS RNA GeoMx experiments. Briefly, after baking for 2 hours at 60° C, sections underwent deparaffinization, heat-induced antigen retrieval (20 minutes at 100°C) with an EDTA based solution (ER2 Leica Cat. AR9640), and proteolytic-induced epitope retrieval (15 minutes at ambient temperature) with proteinase K (Ambion, Cat: AM2546, diluted to 1ug/ml) on a Leica Bond Rx Auto-stainer. After antigen retrieval, slides were moved to 1XPBS until RNA probe hybridization. For whole transcriptome analysis, slides were incubated overnight with 200ul of WTA Mouse RNA probe mix (NanoString®, Cat: 121401102, Lot: MW0030923) consisting of ~18,000 annotated probes with photocleavable tags. Slides were covered with a Hybrislip (Grace biolabs) to ensure uniform distribution of probes and placed in an HybEZ™ oven at 37°C.

Following probe incubation, slides underwent two stringent washes (2XSSC, Formamide 50% v/v, at 37°C) and two 2XSSC (room temperature) washes. Slides were then immune-fluorescently labeled with anti-Troponin I antibody (TnI-Alexa488, Abcam ab196384, clone: EP1106Y, RRID: AB_2910237, Lot: GR213274-7). After slide preparation, slides were loaded onto the instrument for fluorescent scanning at 20X magnification (0.4 µm/px). Regions of interest were then selected based on tissue morphology. 7 to 10 regions of each heart section (including areas from the free wall of either the RV or the LV) were selected for segmentation into TnI(+) and TnI(-) segments that each contained at least 3397 $\mu m^2$. After segmentation, Tag acquisition was performed. The area of tissue within the ROI was precisely illuminated with UV light (~1 $\mu m^2$ resolution) to release the barcodes from the hybridized probes. Each segment was processed individually, and UV-released tags were collected into 96-well microplates. Once all segments were acquired, the multi-well plates were covered and stored at -20C until sequencing at NYUGSM Genome Technology Center.

The sample collection plate was sealed and stored at −20 °C. After thawing, the plates were sealed with AeraSeal film (Excel Scientific), and the aspirates were dried at 65 °C in a thermal cycler with an open top. Once dried, the wells of the collection plate were rehydrated with 10 µl of DEPC water, mixed and spun. A volume of 4 µl each of the DSP aspirate and primer (GeoMx SeqCode Primer plate) were added along with 2 µl of the the 5× PCR Master mix to a new PCR plate and mixed. The sealed plate was transferred to a C1000 thermocycler (Bio-Rad) and Illumina flowcell-compatible libraries were constructed by 18 cycles of PCR amplification according to instructions. Of the resultant libraries (from each well of the PCR plate), 4 µl were pooled into a 1.5 ml tube and cleaned using AMPure XP beads (Beckman Coulter). The quality of the pooled libraries was assessed on tapestation, where the amplicon size was 162 bp and no residual primers were detected. After their quantification on a qPCR, the libraries were sequenced on a NovaSeq6000 SP100 flowcell (Illumina) as paired-end reads with 5% PhiX spike-in. The resultant sequencing reads were demultiplexed using the barcode information provided, and fastq files were generated. The GeoMx NGS pipeline (DND software from nanoString) was then used to generate digital count conversion (DCC) files. The zipped DCC folder was then transferred to DSP and downstream data analysis performed.

The GeoMx NGS gene expression analysis workflow was used for quality control (QC) and normalization. QC was done by filtering segments by low % sequencing saturation and low % aligned reads, followed by principal component analysis (PCA). Threshold for segment filtration was set at 5% or below. To normalize counts, Q3 normalization was performed on the filtered dataset. As part of QC, dimension reduction was conducted using PCA. Remove Unwanted variation (RUV) batch correction was also performed. Normalization was via the limma-voom pipeline (version 3.54.2). [79] Differential expression analysis was conducted using Empirical Bayes (limma version 3.54.2). Analysis was completed using R version 4.2.3. KEGG analysis was performed using KEGGREST version 1.38.0. Datasets acquired from the DISGENET (V24.4) database were of Parkinson’s disease, Alzheimer’s disease, and Amyotrophic lateral sclerosis (ALS). [37]

**Opal 6plx staining (Multiplex immunohistochemistry)**

5 μm sections were immunostained on a Leica BondRx auto-stainer according to the manufacturer's instructions. In brief, sections were deparaffinized online and treated with 3% $H_2O_2$ to inhibit endogenous peroxidases, followed by antigen retrieval with either ER1 (Leica, AR9961; pH6) or ER2 (Leica, AR9640; pH9) retrieval buffer at 100ºC for either 20 or 60 minutes. After blocking, slides were incubated with the first primary antibody and secondary HRP polymer pair. Slides were blocked with Rodent Block M (Biocare, RBM961) then Primary Antibody Diluent (Leica, AR93520) before incubation with primary antibodies raised in rodents, or with only Primary Antibody Diluent before incubation with primary antibodies raised in Rabbit. Following secondary incubation, slides underwent HRP-mediated tyramide signal amplification with a specific Opal® fluorophore. Once the Opal® fluorophore was covalently linked to the antigen, primary and secondary antibodies were removed with a 95ºC retrieval step. This sequence was repeated 5 more times with subsequent primary and secondary antibody pairs, using a different Opal fluorophore with each primary antibody (see **Table S2**). After antibody staining, sections were counterstained with spectral DAPI (Akoya Biosciences, FP1490) and mounted with ProLong Gold Antifade (ThermoFisher Scientific, P36935).

Semi-automated image acquisition was performed on an Akoya Vectra Polaris (PhenoImagerHT) multispectral imaging system. Slides were scanned at 20X magnification using PhenoImagerHT 2.0 software in conjunction with Phenochart 2.0 and InForm 3.0 to generate unmixed whole slide qptiff scans. Image files were uploaded to the NYUGSoM's OMERO Plus image data management system (Glencoe Software).

**Statistical Analysis**

Numeric results are reported as mean ± standard deviation (SD). Data pooled from individual animals were evaluated using the workflow described by Sikkel et al. [20] to assess the validity of the assumption of independence in the presence of potential data clustering. When clustering was detected, hierarchical analysis was performed according to the methods detailed in the supplementary material of Sikkel et al. [20] All datasets were tested for normality using the Shapiro-Wilk test, and statistical significance was determined using either parametric or non-parametric tests, as appropriate. The specific tests used are indicated in the figure legends. Data analysis was conducted using RStudio and GraphPad Prism versions 9.0 and 10.0.
